# Mapping single-cell developmental potential in health and disease with interpretable deep learning

**DOI:** 10.1101/2024.03.19.585637

**Authors:** Minji Kang, Jose Juan Almagro Armenteros, Gunsagar S. Gulati, Rachel Gleyzer, Susanna Avagyan, Erin L. Brown, Wubing Zhang, Abul Usmani, Noah Earland, Zhenqin Wu, James Zou, Ryan C. Fields, David Y. Chen, Aadel A. Chaudhuri, Aaron M. Newman

## Abstract

Single-cell RNA sequencing (scRNA-seq) has transformed our understanding of cell fate in developmental systems. However, identifying the molecular hallmarks of potency – the capacity of a cell to differentiate into other cell types – has remained challenging. Here, we introduce CytoTRACE 2, an interpretable deep learning framework for characterizing potency and differentiation states on an absolute scale from scRNA-seq data. Across 31 human and mouse scRNA-seq datasets encompassing 28 tissue types, CytoTRACE 2 outperformed existing methods for recovering experimentally determined potency levels and differentiation states covering the entire range of cellular ontogeny. Moreover, it reconstructed the temporal hierarchy of mouse embryogenesis across 62 timepoints; identified pan-tissue expression programs that discriminate major potency levels; and facilitated discovery of cellular phenotypes in cancer linked to survival and immunotherapy resistance. Our results illuminate a fundamental feature of cell biology and provide a broadly applicable platform for delineating single-cell differentiation landscapes in health and disease.

## Introduction

All cells, from the fertilized egg to its mature progeny, are hierarchically organized in multicellular life. Such maps of ‘cellular ontogeny’ are exemplified by the lineage tree of *C. elegans*, where every parent-daughter relationship is known^1^. While lineage tracing, functional transplantation assays, and single-cell genomics have revealed key insights into developmental hierarchies^2–5^, the molecular programs underlying potency – a cell’s ability to differentiate into more specialized cell types – remain unclear. For example, totipotent, multipotent, and unipotent cells each have vastly different developmental capacities, yet little is known about the transcriptional profiles that distinguish them. An improved understanding of cell potency would facilitate new insights into diverse physiological and pathological processes, including cancer, where altered differentiation states and plasticity programs shape clinical outcomes^6^.

In recent work, we showed that the number of detectably expressed genes per cell is a hallmark of developmental potential. Leveraging this finding, we developed CytoTRACE, a computational method for predicting cellular maturity from single-cell RNA sequencing (scRNA-seq) data^7^. Despite its broad applicability, CytoTRACE – like other methods for trajectory inference, including scVelo^8^, CellRank^9^, and Monocle 3^10^ – predicts single-cell orderings in a manner that is relative to each dataset (thus, the least differentiated cells in one dataset may have equivalent potency to the most differentiated cells in another). This has made it difficult to unify such predictions across datasets and contextualize them against the backdrop of cellular potency.

To overcome these challenges, we developed CytoTRACE 2, an interpretable deep learning framework for jointly determining single-cell potency categories and absolute developmental potential from scRNA-seq data. Unlike most deep learning methods, which are not inherently explainable, CytoTRACE 2 implements a novel architecture that learns multivariate gene expression programs for each phenotype of interest. Such programs can be seamlessly extracted from the model, are readily interpretable, and deliver remarkably accurate predictions of developmental potential. To demonstrate the utility of CytoTRACE 2, we curated human and mouse scRNA-seq datasets with gold standard potency levels and differentiation trajectories. We then trained, validated, and benchmarked the performance of our approach, explored its potential for reconstructing mouse developmental ontogeny, and used it to uncover candidate differentiation programs in cancer. Our results illuminate pan-tissue determinants of cell potency and highlight the value of interpretable deep learning for characterizing single-cell developmental states in health and disease (https://github.com/digitalcytometry/cytotrace2).

## Results

### Modeling cell potency with interpretable AI

CytoTRACE 2 was designed to provide a unique view of developmental potential. Unlike other methods, it predicts single-cell potency categories and differentiation states on an absolute scale using classically defined developmental stages as “anchor points” (**Methods**). To achieve this goal, we focused on six potency categories spanning the full range of cellular ontogeny: totipotent stem cells capable of generating an entire multicellular organism; pluripotent stem cells with the capacity to differentiate into all adult cell types; lineage-restricted multipotent, oligopotent, and unipotent cells, each capable of producing >3, 2 or 3, or 1 downstream cell type(s), respectively, and differentiated cells, ranging from mature to terminally differentiated phenotypes (**Figure 1A**; additional details in **Methods**). With these annotations, we screened online repositories for scRNA-seq data with assignable potency levels from humans and mice, for which extensive genomic data are available. This yielded a comprehensive training set consisting of 88 cell phenotypes (79 mouse and 20 human), 18 tissue types, 17 datasets, and six platforms, including droplet and plate-based assays, with 176k cells confidently assigned to potency levels in both species (**Figure 1B**, **Methods**).

**Figure 1.**
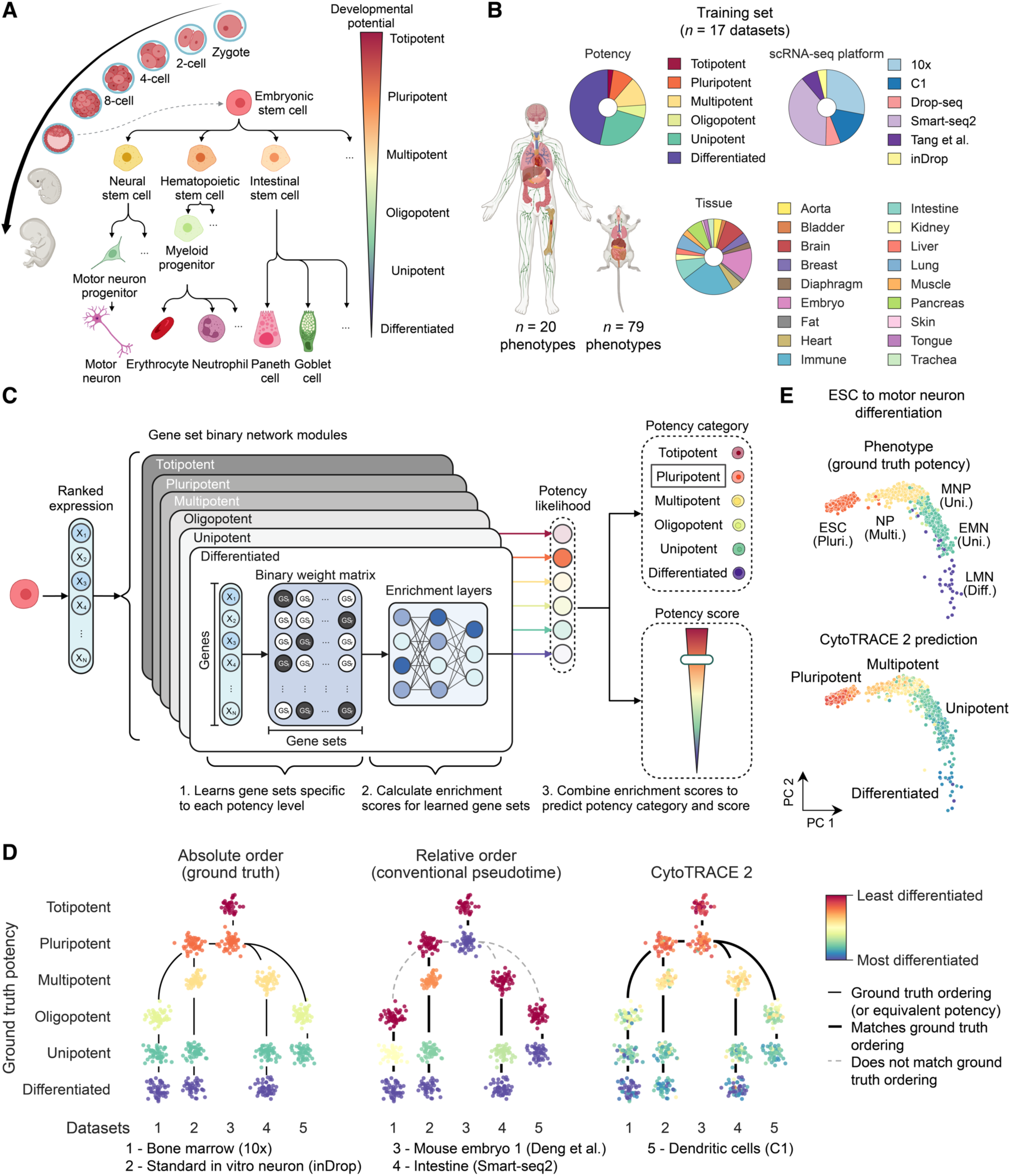
Development of CytoTRACE 2. (**A**) Overview of cell potency. Organized into six major categories, potency describes the capacity of a cell to differentiate into more specialized cell types, beginning with the totipotent zygote and extending through a series of developmental intermediates to fully mature progeny. Details of each potency category are provided in **Methods**. (**B**) Composition of the training set. (**C**) Schematic description of CytoTRACE 2, an interpretable AI system for predicting absolute developmental potential from scRNA-seq data. CytoTRACE 2 employs a multi-step workflow, in which single-cell transcriptomes are first converted to rank space and then processed through a series of neural network components, termed gene set binary network (GSBN) modules, where potency-associated gene sets are decoded (step 1) and single-cell gene set enrichment scores are determined (step 2). Next, potency-specific likelihoods for each cell are converted to categorical and continuous outputs (step 3). The latter, termed a ‘potency score’, ranges from 1 (totipotent) to 0 (differentiated). Additional details are provided in **Figure S1A** and **Methods**. (**D**) *Left:* Illustration of absolute developmental order, in which points denote cells colored by ground truth potency from five independent mouse scRNA-seq datasets (labeled 1 – 5 and organized into columns). Solid lines indicate known lineage relationships, either direct or indirect, and do not imply the absence of developmental intermediates. Phenotypes with >50 cells from a given dataset were uniformly downsampled to 50 cells for clarity. *Center:* Same as the left, but showing conventional pseudotime orderings, or ‘relative order’, in which cells are ordered within a dataset but not across them. *Right:* Same as the left but showing potency scores predicted by CytoTRACE 2. (**E**) PCA embedding of ground truth phenotypes and CytoTRACE 2 potency scores for a dataset covering embryonic stem cell differentiation into motor neurons (*n* = 1,726 cells; ‘Standard in vitro neuron, inDrop’ dataset^61^). ESC, embryonic stem cell; NP, neural progenitor; MNP, motor neuron progenitor; EMN, early motor neuron; LMN, late motor neuron. A – C were created using icons from BioRender.com. See also **Figure S1**.

Like its predecessor, we developed CytoTRACE 2 with an emphasis on understanding the molecular determinants of developmental potential. However, since most deep learning methods require dedicated procedures to ascertain feature importance^11,12^, we designed CytoTRACE 2 with a novel, fully explainable architecture for single-cell classification tasks (**Methods**). Drawing inspiration from binarized neural networks^13^, where weights take values of 1 or –1, our approach, termed a Gene Set Binary Network (GSBN), automatically activates (=1) or deactivates (=0) individual genes, with the goal of identifying highly discriminative gene sets that best describe each phenotype of interest (e.g., potency levels). Multiple gene sets can be learned per phenotype, enhancing flexibility, and informative genes can be easily extracted from the model. As such, unlike the majority of deep learning architectures, GSBNs can yield immediate insight into the molecular programs underpinning performance (**Methods**).

Leveraging this approach to model all six potency categories, CytoTRACE 2 proceeds in five main steps: input standardization, discovery and recovery of potency-specific molecular programs (during and after training, respectively), determination of single-cell enrichment scores for each molecular program, and aggregation of single-cell enrichment scores into two outputs for each cell: (1) the probability of each potency category and (2) a continuous potency score – ranging from 1 (totipotent) to 0 (differentiated) (**Figures 1C**, **S1A**, and **S1B**; **Methods**). Based on the assumption that transcriptionally similar cells occupy related differentiation states, in the fifth step, CytoTRACE 2 leverages a nearest neighbor approach to smooth individual differentiation scores (**Figures S1C** and **S1D**). Additional details are provided in **Methods.**

### Technical performance and benchmarking of CytoTRACE 2

Having established a large compendium of training data, we next assessed the performance of CytoTRACE 2. For completeness, we quantified accuracy using two definitions of developmental ordering: ‘absolute order’, in which predictions from all datasets were analyzed in aggregate with respect to known potency categories, and ‘relative order’, in which cells of each dataset were evaluated on a relative scale – from the least to most differentiated (**Figure 1D**, left and center). With the latter, we could also evaluate differences between granular differentiation states assigned to the same potency category (e.g., multipotent hematopoietic and intestinal adult stem cells versus their immediate downstream multipotent progenitors). For both measures, we quantified the agreement between known and predicted developmental orderings using weighted Kendall correlation. This allowed us to assess concordance between ranks while balancing key variables to minimize bias (**Methods**).

We began by optimizing model hyperparameters using a Bayesian cross-validation scheme (**Methods**). Across a wide range of values, we observed only minimal variation in performance, with a slight preference for >5 gene sets per potency category (**Figure S1B**). As such, we selected robust hyperparameter values to retrain the model and employed leave-one-out cross-validation as a first step toward evaluating it (**Methods**).

On the training set, CytoTRACE 2 achieved remarkably high accuracy, immediately distinguishing it from pseudotime methods that infer relative orderings (**Figures 1D, 1E, 2A**, and **2B**). This was true regardless of whether we assessed results at the single-cell level or when aggregating by phenotype (**Figures 2A** and **2B**). Moreover, on a continuous scale ranging from 1 (totipotent) to 0 (differentiated), where each potency category was uniformly distributed, 90% of cells were assigned within 0.1 units of the correct potency label (**Figure S2A**). This result was highly statistically significant (*P* < 10^−6^; **Figure S2A**), underscoring the promise of CytoTRACE 2 for determining single-cell developmental states in an absolute manner.

**Figure 2.**
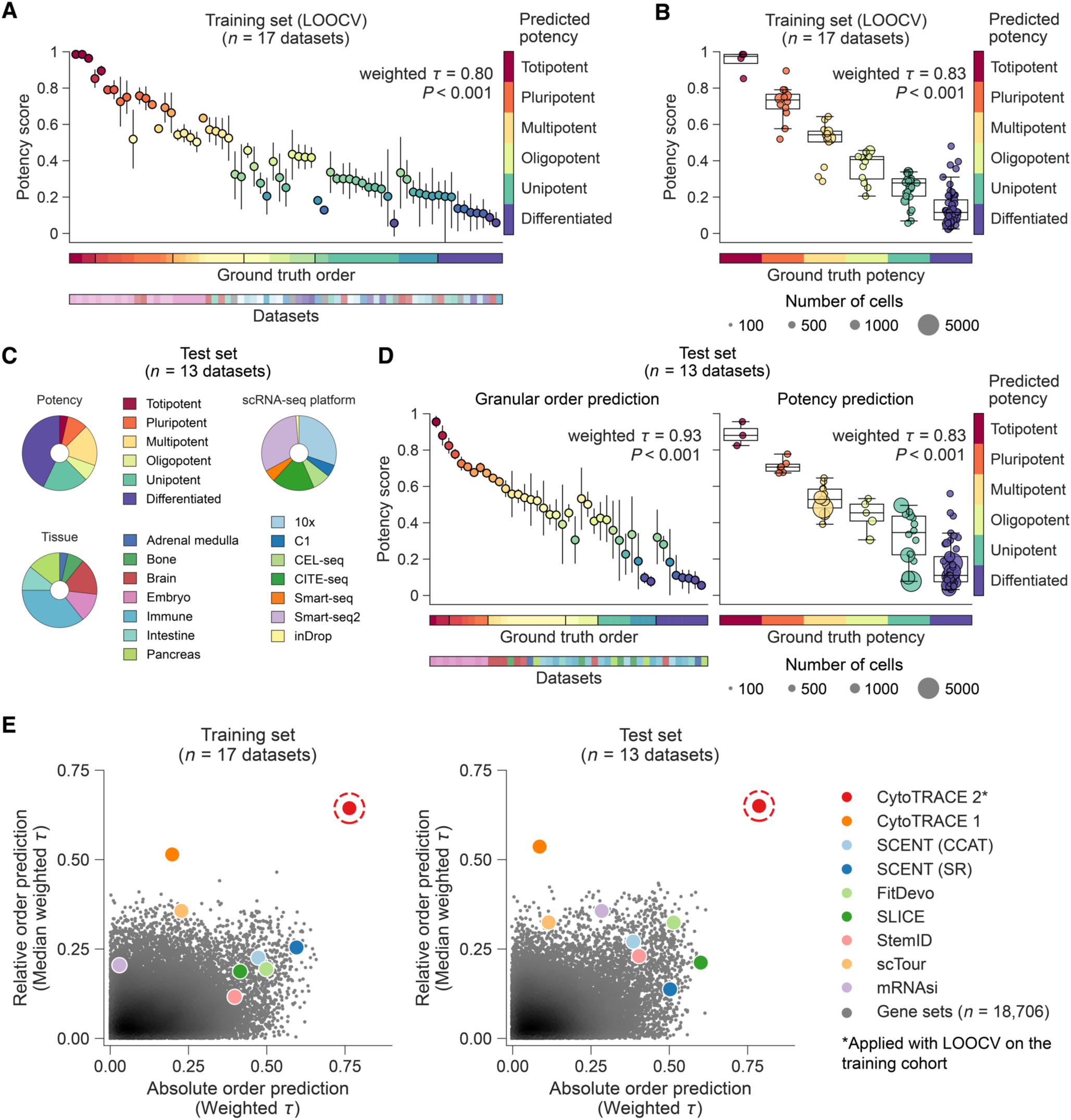
Technical performance of CytoTRACE 2. (**A**) Performance of CytoTRACE 2 on the training set using leave-one-dataset-out cross-validation (LOOCV). Shown are continuous and categorical potency levels predicted by CytoTRACE 2 (left and right y-axes, respectively) versus ground truth developmental states (*n* = 23; x-axis). The latter combines six major potency categories (Figure 1A) with more granular developmental states described in **Methods**. The position and color of each data point reflect the mean CytoTRACE 2 potency score for a given dataset stratified by ground truth. For points with multiple phenotypes, each phenotype is represented equally. Error bars represent 95% CI. Concordance with ground truth was determined using Kendall correlation weighted as described in **Methods**, with statistical significance determined asymptotically via two-sided Z-test. For visualization, points are arranged by decreasing potency score within each ground truth state. (**B**) Same as A but showing box plots of the mean CytoTRACE 2 score for each phenotype / dataset pair, stratified by ground truth potency (center line, median; box limits, upper and lower quartiles; whiskers, largest and smallest values within 1.5 × IQR of the box limits; IQR, interquartile range). Performance was determined as in A but relative to six potency categories as described in **Methods.** Circle size is proportional to the number of cells analyzed. (**C**) Composition of the held-out test set. (**D**) Same as A and B but shown for the test set. Here, CytoTRACE 2 results were ensemble-averaged across 17 models learned on the training set (**Methods**). (**E**) Scatter plots comparing CytoTRACE 2 potency scores against eight previous methods and 18,706 annotated gene sets for determining developmental orderings in the training (left) and test (right) sets. Results are quantified using weighted Kendall correlation, applied to evaluate absolute order (cross-dataset correlation across six potency categories, x-axis) and relative order (within-dataset correlation, y-axis) (see also Figure 1D; **Methods**). For clarity, the absolute value of each metric is shown. All comparisons were assessed at single-cell resolution; weighting schemes and raw data are provided in **Methods**. The asterisk denotes the use of LOOCV for CytoTRACE 2 on the training set. Among gene sets, darker gray denotes higher point density. See also **Figures S2** and **S3**.

To evaluate generalizability, we next extended our analysis to fully unseen data. To do so, we assembled a held-out test set comprised of 13 additional human and mouse scRNA-seq datasets encompassing experimentally confirmed differentiation trajectories, six potency categories, seven tissue systems, seven platforms, and 363k evaluable cells (**Figure 2C**). Across diverse metrics, CytoTRACE 2 exhibited nearly identical performance to the training set and was not significantly affected by differences in species, tissue type, or platform (**Figures 2D** and **S2B** to **S2D**). Moreover, we observed no significant loss of performance when analyzing phenotypes and tissue types that were absent during model training (**Figure S2E**). Thus, CytoTRACE 2 can learn molecular profiles of cell potency that generalize to new datasets, including previously unseen phenotypic states.

We next asked whether CytoTRACE 2 is robust to dropout and variation in the number of sequenced RNA molecules per cell. Indeed, by perturbing the number and diversity of unique molecular identifiers (UMIs) per cell from droplet-based datasets in the test set, we found that estimates of developmental potential were virtually identical down to 1,000 UMIs per cell, with only modest effects observed at 750 UMIs per cell (**Figure S2F**, top). Similar results were obtained when down-sampling UMIs by a fixed percentage (**Figure S2F**, bottom). Consistent with this, when analyzing developmental systems in which the known ordering was inverted by the original CytoTRACE^7^ owing to the presence of immature cells with low mRNA content, CytoTRACE 2 corrected the inversion (**Figures S2G** and **S2H**). Thus, unlike its predecessor^7^, sparsity and mRNA content are not major factors influencing performance when practically obtainable count thresholds are met.

Given these favorable results, we next benchmarked CytoTRACE 2 against competing measures of single-cell developmental potential. To promote a comprehensive assessment, we focused on measures with broad applicability to developmental systems independent of the number of transitional states or the presence of specific developmental timescales (**Methods**). In total, we included eight dedicated methods based on gene counts (CytoTRACE 1^7^), transcriptional entropy (SCENT CCAT^14^, SCENT SR^15^, StemID2^16^, SLICE^17^), and machine learning (mRNAsi^18^, scTour^19^, FitDevo^20^), and 18,706 annotated gene sets, consisting of transcription factor binding targets^21,22^ and the entire Molecular Signatures Database^23^ (**Methods**). To quantify performance on the training and test sets, we determined absolute and relative orderings as above. Regardless of the cohort or the definition of developmental ordering, CytoTRACE 2 emerged as the leading approach, outperforming all evaluated methods and gene signatures, in most cases, by a substantial margin (**Figure 2E**).

To further authenticate performance, we next extended our analysis to Tabula Sapiens^24^, a multi-donor scRNA-seq atlas covering 483,152 postmortem cells from 22 human tissue types. We also evaluated CytoTRACE 2 against pseudotime trajectories determined by RNA velocity as implemented by scVelo^8^. Applied to Tabula Sapiens, CytoTRACE 2 was more accurate than previous methods (**Figure S3A**). Likewise, CytoTRACE 2 exhibited favorable or superior performance when compared against the RNA velocity kernel in scVelo while also providing estimates of absolute developmental potential (**Figures S3B** to **S3D**).

Together, these results validate CytoTRACE 2, establish its extensibility to new data, and emphasize its potential for revealing differentiation states and biological insights that are unattainable by existing in silico methods and gene signatures.

### Tracing developmental potential from the zygote to birth

In recent years, tissue and organ-level cell atlases have generated millions of single-cell expression profiles^25^, however many biological systems are only partially covered by individual datasets. Given that CytoTRACE 2 is anchored to a predefined developmental range, we sought to further test its ability to overcome technical variation between scRNA-seq datasets and achieve large-scale integrative analysis of cell potency.

We began by asking whether dataset-specific technical variation has any discernible effect on CytoTRACE 2. To this end, we examined mouse bone marrow hematopoiesis, a well-established developmental system covered by two datasets in the scRNA-seq cohorts described above. Using fastMNN^26^ to integrate the scRNA-seq expression matrices, CytoTRACE 2 predictions were comparably accurate before and after technical variation was removed (**Figures S3E** and **S3F**). This was consistent with our earlier benchmarking results, in which strong inter-dataset performance was observed despite running CytoTRACE 2 on each dataset separately (**Figure 2E**).

We next turned to mouse embryogenesis, with the goal of charting single-cell potency over the full prenatal period. Notably, no existing atlas covers this entire process at daily intervals, providing an opportunity to evaluate the consistency of potency predictions across multiple datasets without batch correction. To achieve continuity, we assembled six timelapse datasets consisting of >11M mouse single-cell transcriptomes^4,27–31^, 62 developmental time points, and five platforms, including traditional plate-based methods, droplet sequencing, and combinatorial index sequencing of single-cell nuclei. Together, these longitudinal data encompass every embryonic day from E0.5 (zygote) through P0 (birth) (**Figure 3A**, bottom). We then applied CytoTRACE 2 to each dataset individually and aggregated the resulting potency scores, both by author-supplied phenotypes and major embryonic time points (**Figure 3A**). Mean potency scores produced by CytoTRACE 2 were strikingly well-correlated with developmental time (absolute Kendall correlation = 0.95, *P* = 1.2 × 10^−24^, **Figure 3B**). Moreover, as expected, overall potency levels tracked with ontogenesis in a graded fashion, with CytoTRACE 2 documenting the transition from totipotent to pluripotent cells during pre-gastrulation; from pluripotent to multipotent cells during gastrulation; and from multipotent to differentiated states during organogenesis (**Figure 3A**).

**Figure 3.**
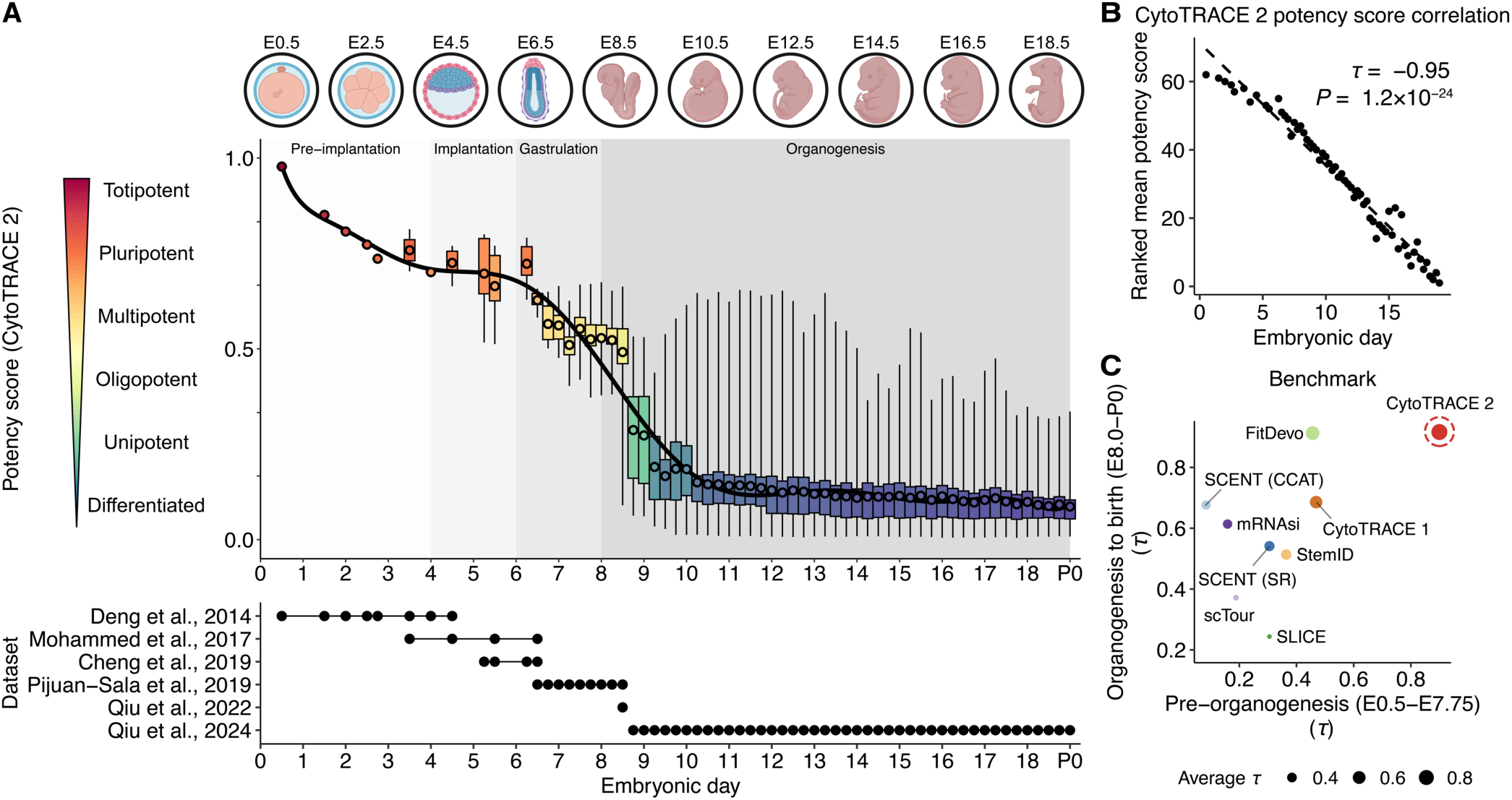
Temporal reconstruction of developmental potential during mouse embryogenesis. (**A**) *Top:* Potency scores determined by CytoTRACE 2 across 62 developmental time points spanning mouse embryogenesis, from the zygote (E0.5) to birth (P0). Four main phases of embryogenesis are indicated: pre-implantation, implantation, gastrulation, and organogenesis^62,63^. Box plots summarizing the average single-cell potency score for each time point and author-supplied phenotype (center circle, mean; box limits, upper and lower quartiles; whiskers, maximum and minimum; IQR, interquartile range). Polynomial regression (6th degree) was applied to visualize the relationship between mean potency scores and time points (solid line). *Bottom*: Overview of six scRNA-seq datasets analyzed by CytoTRACE 2 and the time points profiled by each. To benchmark CytoTRACE 2 against methods with highly variable efficiencies (panel C), cells from each dataset were downsampled as described in **Methods**, resulting in 158,025 evaluable cells. (**B**) Linearity between average potency scores expressed in rank space (y-axis) and the corresponding time points (*n* = 62; x-axis). Concordance was calculated using linear regression (dashed line) and Kendall correlation (𝜏), with the latter weighted by the number of time points per embryonic day. Statistical significance was determined using a two-sided Z-test. (**C**) Scatter plot comparing the performance of CytoTRACE 2 to previous approaches (eight methods) for reconstructing the temporal hierarchy of 17 time points preceding organogenesis (y-axis) versus 45 time points spanning organogenesis (beginning at E8.0^63^) to birth (x-axis). For each approach, points represent the Kendall correlation between the indicated time points and the average prediction per time point, with the latter represented in rank space. Correlations are weighted by whole day intervals to account for imbalances in the number of evaluable time points per day. Point sizes represent the average weighted Kendall correlation per approach. See **Methods** for additional details. The top of A was created using icons from BioRender.com.

To benchmark this result, we applied eight previous methods to each of these datasets (**Methods**). Regardless of how we analyzed concordance with developmental time, CytoTRACE 2 showed performance gains over competing methods, including CytoTRACE 1 (**Figure 3C**). This was evident before and during organogenesis where both dramatic and subtler changes in median potency levels were respectively observed (**Figure 3C**). These data further demonstrate the potential of CytoTRACE 2 to harmonize predictions across scRNA-seq datasets, experimental batches, and distinct protocols, highlighting its potential for single-cell potency assessment at massive scale.

### Potency-related programs determined by interpretable AI

Having evaluated the technical performance of CytoTRACE 2, we next leveraged its inherent interpretability to explore the molecular correlates of developmental potential (**Figure 4A**). Notably, previous genomic surveys of stem cell transcriptional programs have largely focused on molecular features of pluripotent stem cells arising from the inner cell mass or somatic cell reprogramming^18,32,33^. In comparison, relatively few studies have examined conserved gene expression signatures of other potency levels^34,35^, and none have characterized potency-specific transcriptional programs at the granularity analyzed in this work.

**Figure 4.**
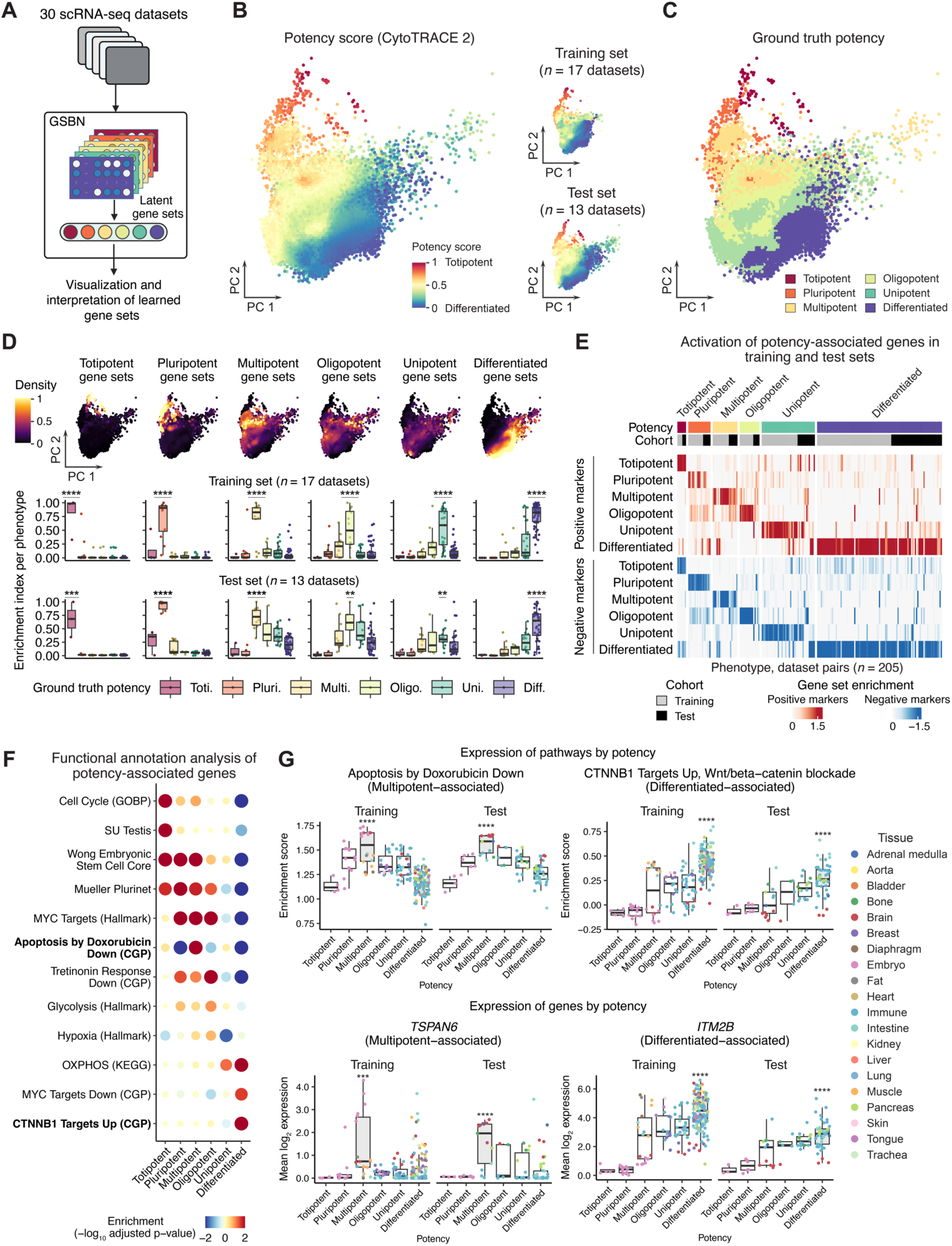
Pan-tissue potency programs determined by interpretable AI. (**A–G**) Visualization and interpretation of potency-associated molecular programs learned by CytoTRACE 2. (**A**) Schematic for characterizing latent gene sets and their activity levels in the training and test sets (*n* = 30 scRNA-seq datasets). GSBN, gene set binary network (see also Figures 1C and **S1A**). (**B**) *Left:* PCA visualization of gene set expression levels in the training and test sets. To mitigate variation in cell density, results for individual cells are aggregated into consecutive windows of 1.5 PC units squared. Points represent individual windows colored by the average CytoTRACE 2 potency score (**Methods**). *Right:* Same as the left panel but stratified by training and test sets. (**C**) Same as B (left) but colored by ground truth potency. (**D**) *Top:* Same as B (left) but colored by the joint enrichment of potency-associated gene sets in each single cell, termed an enrichment index (**Methods**). *Center:* Box plot showing the mean enrichment index of potency-associated gene sets in the training set. Cells are organized by ground truth potency and aggregated by cell type (within each dataset). *Bottom:* Same as the center panel but shown for the test set. (**E**) Similar to D but depicted as a heat map in which enrichment scores from datasets with at least two ground truth potency categories are dichotomized by gene set-specific weights learned by CytoTRACE 2 and averaged by potency category (positive weights, top; negative weights, bottom). Positive and negative weights select for positive and negative markers of cell potency, respectively. Enrichment scores were calculated and normalized as described in **Methods**. (**F**) Bubble plot showing functional annotation of potency-associated genes learned by CytoTRACE 2. Enrichments were determined using pre-ranked GSEA applied to 15,634 molecular signatures. Selected results are highlighted, with bolded entries further analyzed in G. (**G**) *Top left:* Box plot showing single-sample GSEA^64^ (ssGSEA) enrichment scores of leading-edge genes within the GRAESSMANN_APOPTOSIS_BY_DOXORUBICIN_DN signature, the most specific signature for positive markers of multipotency learned by CytoTRACE 2. Expression data were first averaged by cell type within each dataset (with the exception of Tabula Muris, which was further divided by tissue / platform pairs), then analyzed by ssGSEA in training and test sets. Data points are colored by tissue type. *Top right:* Same as the top left panel but for FEVR_CTNNB1_TARGETS_UP, a signature with high specificity for differentiated cells. *Bottom:* Same as the top panel but showing the expression of single genes with maximal specificity for multipotent and differentiated cells. Genes and gene sets were selected as described in **Methods**. In D and G, the box center lines, bounds of the box, and whiskers denote medians, 1st and 3rd quartiles, and minimum and maximum values within 1.5 × IQR (interquartile range) of the box limits, respectively. Permutation tests were applied to determine statistical significance in D and G (**Methods**). \*\**P* < 0.01; \*\*\**P* < 0.001; \*\*\*\**P* < 0.0001. Panel A was created using BioRender.com. See also **Figure S4**.

Therefore, to bridge this gap, we began by visualizing CytoTRACE 2’s internal representation of gene expression signatures (**Figure 4A**). We first compiled results from our earlier benchmarking analysis of 30 training and test datasets (**Figure 2**). We then converted gene set activity levels from the GSBN modules into a low dimensional embedding using principal components analysis (PCA) (**Figure 4B**). Remarkably, despite combining nearly 100k cells from 30 scRNA-seq datasets, 9 platforms, and 20 tissue types, predicted differentiation states formed a cohesive gradient spanning the breadth of cellular ontogeny (**Figure 4B**, left). This gradient was independent of training or test sets (**Figure 4B**, right), faithful to ground truth potency categories (**Figure 4C**), and independent of obvious batch effects (**Figure S4A**). Moreover, gene set activities from each potency category exhibited a banding pattern marking consecutive potency levels (**Figure 4D**, top). This result was confirmed quantitatively in training and test sets (**Figure 4D**, middle and bottom), emphasizing both the discriminatory power of the gene sets learned by the model and the utility of our GSBN approach.

To dissect the biology underlying these programs, we next interrogated their molecular themes. As part of this process, we dichotomized gene sets based on their polarity in the model (**Figure S4B**). We found that gene sets with positive weights were strongly enriched in a given potency category, whereas those with negative weights were preferentially depleted (**Figure 4E**). To capture each gene’s importance and directionality in the model, we created six ranked gene lists, one for each potency level (**Methods**). We then applied gene set enrichment analysis (GSEA) to annotate these lists, revealing both expected and unexpected functional themes (**Methods**). For example, many stem cells inhabit niches with lower oxygen concentrations and rely on anerobic glycolysis for energy production^36^. Conversely, more differentiated phenotypes generally shift their metabolic requirements toward oxidative phosphorylation^36^. These trends, along with upregulation of canonical stem cell programs in the least mature cells (e.g., expression of MYC target genes^37^), were confirmed by GSEA (**Figure 4F**). Moreover, as expected, embryonic stem cell signatures were highly discriminatory for pluripotency, with *POU5F1*, a canonical pluripotency gene encoding OCT4^38,39^, ranking in the top 0.3% of genes (**Figure 4F**). Additionally, chromosome organization and cell cycle processes were most enriched in totipotency, consistent with the rapid nuclear division and elevated metabolism that accompany the earliest events of embryogenesis^40^ (**Figure 4F**).

We also identified molecular signatures with unexpectedly robust specificity for potency levels in tissue-resident cells (**Figure 4F**). However, many of these signatures were derived from perturbation experiments rather than canonical pathways, illuminating ill-defined biology. For example, genes downregulated by doxorubicin in ME-A breast cancer cells^41^ were most specific for multipotency (**Figure S4G**, top left). Within this signature, we discovered a unique lipid metabolism network with specificity for multipotent cells across diverse tissue types (**Figures S4C** to **S4E; Methods**). Similarly, CytoTRACE 2 genes associated with oligopotent and differentiated cells were preferentially enriched in perturbation programs, including genes regulated by exposure to all-trans retinoic acid^42^ (**Figure S4F**) and blockade of Wnt/beta-catenin signaling^43^ (**Figure 4G**, top right), respectively. These trends were confirmed in held-out test data (**Figures S4F** and **4G**, top), both corroborating our findings and revealing previously unknown expression programs of cell potency with pan-tissue representation.

In line with these data, several potency-associated genes were shared across distinct cellular phenotypes and developmental origins. For example, among the top 50 genes most highly ranked by the model in each potency category, *TSPAN6* and *ITM2B* each showed multi-tissue conservation as positive markers of multipotent and differentiated cells, respectively (**Figures 4G**, bottom, and **S4G**; **Methods**).

Thus, owing to the interpretability, expressiveness, and performance characteristics of CytoTRACE 2, we could readily identify both known and novel molecular correlates of absolute differentiation status in humans and mice. To facilitate further analysis by others, functional enrichment data, leading edge genes, and putative transcription factors are provided in the supplement.

### Single-cell developmental states across human malignancies

In cancer, stem cell regulatory programs are strongly implicated in tumorigenesis, therapy resistance, immune evasion, and metastasis^44–47^. However, deciphering cancer cell differentiation states with single-cell genomics has remained challenging, especially when combining results across different samples, tumor types, and platforms. Therefore, to further explore the utility of CytoTRACE 2, we used it to identify candidate differentiation states on an absolute scale across human malignancies.

We began by asking whether expression signatures of diverse potency categories derived from normal developmental systems portend survival outcomes. Indeed, when analyzing gene-level survival associations across 39 human malignancies from PRECOG^48^, signatures of increasingly immature potency levels learned by CytoTRACE 2, excluding totipotency, were correlated with shorter survival time (*Q* < 0.05; **Figure S5A**, left). Moreover, when omitting genes linked to pluripotent stem cells and proliferation, this trend persisted in signatures of multipotent cells and their progeny (**Figure S5A**, right). Thus, despite the widespread use of pluripotency signatures as surrogates of cellular differentiation status^18,49–51^, these results reinforce the notion that classically defined potency categories are fundamentally distinct.

To expand upon this finding, we applied CytoTRACE 2 to 458,226 previously annotated malignant cells from 17 cancer types and 46 scRNA-seq datasets covering 591 patients with carcinoma, melanoma, sarcoma, blood cancer, or brain cancer^52^ (**Figure 5A**, left). When median-aggregated across cancer types, over half of malignant cells were predicted to be differentiated (60%), with unipotent-like, oligopotent-like, and multipotent-like cells each encompassing 3%, 21%, and 9% of malignant cells, respectively. Consistent with expectation, pluripotent-like and totipotent-like phenotypes were either extremely rare (0.18% on average) or not detected, respectively, consistent with the absence of teratomas in these samples (**Figure S5B**). For acute myeloid leukemia (AML), a positive control in which cellular hierarchies are established, markers of predicted potency levels were well-validated by published signatures of leukemic stem cells and their progeny^53^ (**Figure S5C**). Moreover, such classifications were largely independent of inferred copy number profiles, in line with epigenetic alterations as a major source of phenotypic variation^6,52,54^ (**Figures S5D** and **S5E**).

**Figure 5.**
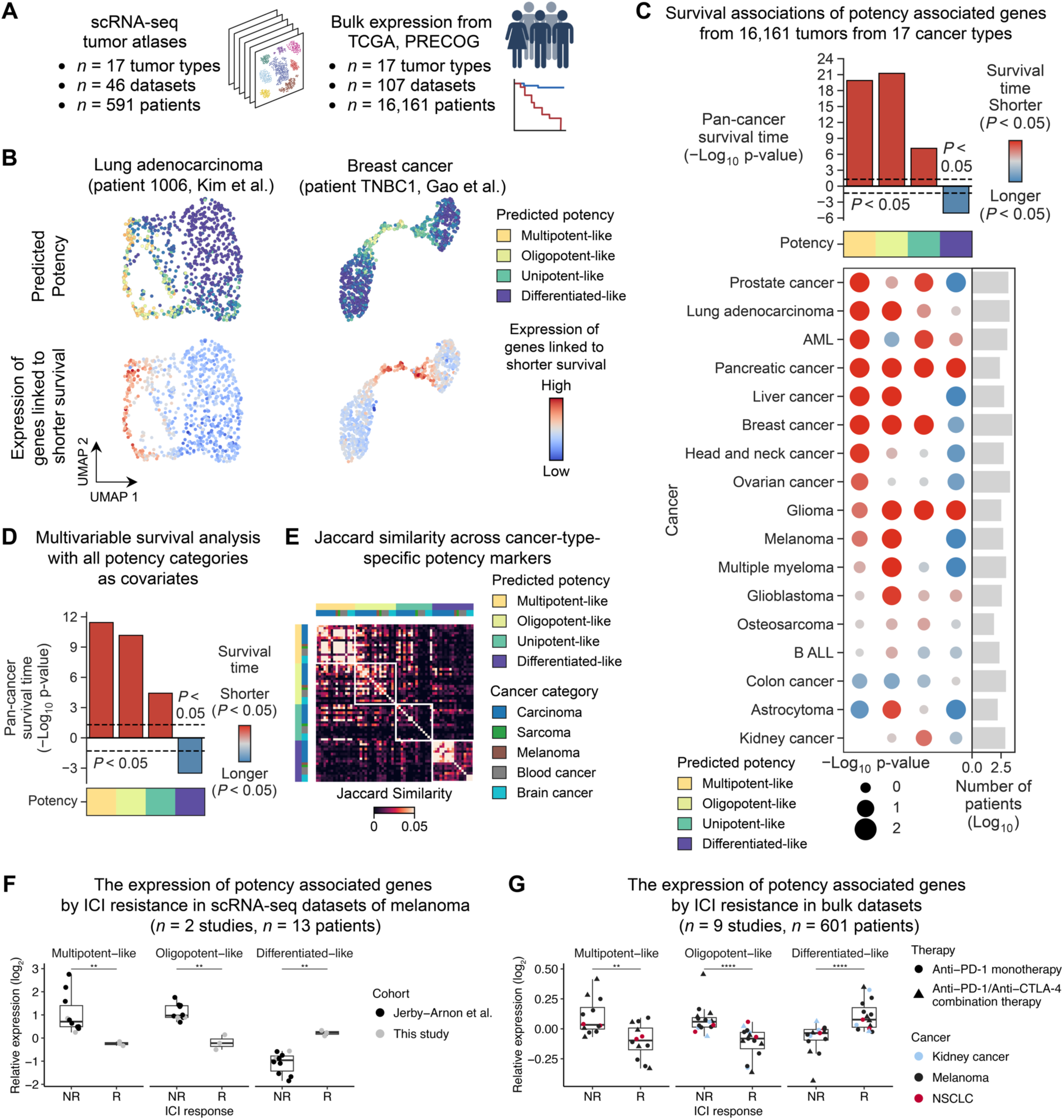
Application of CytoTRACE 2 to neoplastic disease. (**A–E**) Multimodal survey of candidate developmental programs (multipotent-like, oligopotent-like, etc.) in 458k malignant cells and 16.2k bulk tumors across 17 human cancer types. (**A**) Inventory of datasets analyzed in A–E. (**B**) UMAP embeddings of malignant cells in representative tumor specimens from patients with lung adenocarcinoma (left) and breast cancer (right). Plots are colored by CytoTRACE 2 potency scores (top) and differential enrichment of the top 1,000 most adversely prognostic genes from bulk tumors, matched by cancer type (bottom) (**Methods**). (**C**) Bubble plot showing univariable survival associations of potency-specific markers in bulk tumor expression profiles from 17 cancer types. Survival associations were integrated across datasets via weighted meta-Z statistics and expressed as –log_10_ p-values using bubble color and size. Pan-cancer survival associations for each potency category (weighted equally by cancer type) are shown above. (**D**) Same as C (top) but after multivariable adjustment for all potency categories detected in each cancer type. (**E**) Heat map depicting the overlap of potency markers predicted for each cancer type and potency category. Similarity was calculated using the Jaccard Index for the top 200 markers identified for each potency category within each cancer type. (**F**, **G**) Association between candidate developmental programs and resistance to immune checkpoint inhibitors (ICIs). (**F**) Box plot showing the mean log_2_ expression of potency-specific markers in melanoma cells profiled by scRNA-seq, organized by ICI response in 13 patients from two datasets and normalized as described in **Methods**. NR, non-responder (*n* = 10 patients); R, responder (*n* = 3 patients). (**G**) Box plot showing the mean log_2_ expression of potency-specific markers averaged by response status and dataset in bulk tumor expression profiles from three cancer types and 601 patients treated with single or combination ICIs (*n* = 9 studies). Data reflect a combination of pre-, on-, and post-treatment tumors. Data were normalized as described in **Methods**. NSCLC, non-small cell lung cancer. In F and G, the box center lines, bounds of the box, and whiskers denote medians, 1st and 3rd quartiles, and minimum and maximum values within 1.5 × IQR (interquartile range) of the box limits, respectively. Statistical significance in F and G was determined using a two-sided Wilcoxon test. \*\**P* < 0.01; \*\*\*\**P* < 0.0001. See also **Figure S5**.

To determine whether these predictions are clinically relevant, we next collected 16.2k bulk tumor expression profiles from matching cancer types with overall survival data from The Cancer Genome Atlas (TCGA)^55^ and PRECOG^48^ (**Figure 5A**, right; **Methods**). After calculating gene-level survival associations for each cancer type, we observed a striking correspondence between malignant cells with higher predicted potency levels and those with elevated expression of genes linked to inferior outcomes (**Figure 5B**). To substantiate this finding, we used the single-cell potency predictions from CytoTRACE 2 to define potency-associated markers for each cancer type and queried their expression in bulk tumor transcriptomes from the same malignancies (**Methods**). Again, we observed highly significant survival associations across malignancies, with elevated multipotent- and oligopotent-like signatures forecasting the poorest outcomes (**Figure 5C**). Importantly, such signatures were more prognostic than those learned from normal developmental systems, highlighting the value of defining potency-associated cellular states de novo with CytoTRACE 2.

We next assessed cancer-specific potency profiles using multivariable analysis. When evaluated across tumor types, the majority of potency profiles were independently prognostic, both with respect to each other (**Figures 5D** and **5E**) and in relation to proliferation genes, pluripotency signatures, stage, age, sex, tumor purity, tumor grade, mRNAsi^18^ (a machine learning-based pluripotency score), and TmS, a recently introduced measure of tumor RNA content^56^. Other programs typically elevated in cancer, such as epithelial mesenchymal transition (EMT), MYC-related processes, and recently described pan-cancer expression modules^52^, were also insufficient to fully explain our findings, pointing toward distinct biology (**Figure S5G**).

Given these results, along with increasing evidence that stem cell and plasticity programs contribute to drug resistance^6,46,57^, we reasoned that potency markers might associate with therapy response. As a step toward addressing this, we explored several preliminary scenarios involving liquid and solid malignancies, multiple therapy types, and in vitro and in vivo settings. For example, in AML, a previously described 7-gene signature linked to cellular immaturity and differential response to 105 drugs (LinClass-7) was most enriched in multipotent-like AML cells, consistent with expectation^53^ (**Figure S5H**). Moreover, in scRNA-seq profiles of melanoma, including new data generated in this work (**Methods**), signatures of immature malignant cells identified by CytoTRACE 2 were significantly higher in tumors with both previous and future resistance to immune checkpoint inhibitors (ICIs) (**Figure 5F**). These findings were recapitulated in a pan-cancer analysis of bulk RNA-seq data covering six cancer types, including 601 pre- and post-treatment human tumors (**Figure 5G**) and 135 mouse tumor models (**Figure S5I**), treated with either single or combination ICI therapy.

Collectively, these data further validate CytoTRACE 2, demonstrate its ability to identify candidate differentiation states from poorly understood developmental systems, and provide a platform for the identification of novel diagnostics and therapeutic targets.

## Discussion

In this study, we describe CytoTRACE 2 as an interpretable deep learning framework for predicting classically defined cell potency labels and differentiation states on an absolute scale from scRNA-seq data. CytoTRACE 2 is the first in silico method designed to achieve both goals, while providing an explicit readout of the molecular profiles that drive performance.

Previous methods for profiling differentiation states from scRNA-seq data are limited by technical biases and cross-dataset interpretability, with no mechanism to explicitly link predictions of stemness/pseudotime to absolute developmental potential^7–10,14–20,58^. Separately, most deep learning architectures are not inherently explainable and require indirect strategies to extract feature importance^11,12^. Leveraging a modular architecture that achieves full transparency, we trained and validated CytoTRACE 2 on a rich repertoire of tissue types, organs, and developmental systems, spanning all major potency levels in humans and mice. With these data, we identified conserved signatures of cell potency and established a resource and analytical tool to dissect developmental potential with high granularity from existing and emerging cell atlases.

We anticipate that CytoTRACE 2 will facilitate important applications beyond those demonstrated in this work. For example, it can be applied to (i) optimize and troubleshoot somatic cell reprogramming and stem cell engineering experiments where the generation of cells with specific potency levels is desired; (ii) distinguish developmental variation from non-developmental state changes in scRNA-seq data; and (iii) promote the discovery of developmental phenotypes and associated tissue microenvironments from scRNA-seq data integrated with spatial transcriptomics data. Importantly, the analytical principles underlying our GSBN approach are completely general, distinct from methods that factorize scRNA-seq data by known gene sets^59,60^ (**Methods**), and should be extensible to any organism, phenotypic state, or data type for which suitable training data are available. Future work will be needed to formally explore these possibilities.

In summary, CytoTRACE 2 is an interpretable AI framework for cell potency assessment from scRNA-seq data. Given its unique capabilities, it should have immediate utility for gaining insights into the cellular determinants of tissue formation and function, and for empowering basic and clinical research into diseases where altered developmental hierarchies play a role.

### Limitations of the study

This study has several limitations. First, like all supervised machine learning strategies, CytoTRACE 2 is dependent upon the quality and breadth of the training data, the flexibility and robustness of the model, and the generalizability of the signal. Aspects of developmental potential that are insufficiently captured by the current model may be addressable with modifications to the training set, the GSBN architecture, or both. Second, while CytoTRACE 2 is generally insensitive to variation in gene/UMI counts per cell, additional optimization will be required for cells with exceedingly low counts (<500 UMIs). Third, ortholog mapping may allow the current model to generalize to species beyond humans and mice but will require additional testing. Finally, the differentiation states and molecular signatures predicted in this work are hypothesis-generating and will require independent confirmation and functional validation in future studies.

## Author contributions

A.M.N., M.K., J.J.A.A., and G.S.G. conceived of the study, developed strategies for related experiments, and wrote the paper with assistance from E.L.B. and R.G. J.J.A.A., M.K., and A.M.N. developed and implemented CytoTRACE 2 with assistance from S.A. and E.L.B. G.S.G. determined potency annotations with assistance from M.K. and A.M.N. M.K., J.J.A.A., and A.M.N. performed data analysis and interpretation with assistance from R.G., G.S.G., S.A., and W.Z. Z.W. assisted with benchmarking under the supervision of J.Z. R.C.F. and D.Y.C. procured melanoma tumor specimens. A.U., N.E., D.Y.C., and A.A.C. performed single-cell expression profiling. All authors commented on the manuscript at all stages.

## Acknowledgments

The authors would like to acknowledge M. Clarke, A. Gentles, J. D’Silva, and S. Sikandar for providing critical feedback on this manuscript. We are grateful to S. Thapa for assistance with software testing. This work was supported by a Stanford Bio-X Interdisciplinary Graduate Fellowship (M.K.), a Stanford Dean’s Postdoctoral Fellowship (J.J.A.A.), the National Science Foundation (R.G., Graduate Research Fellowship), donor funds from the Barnes-Jewish Foundation (D.Y.C.), the V Foundation for Cancer Research V Scholar Award (A.A.C.), the Melanoma Research Alliance (D.Y.C., A.A.C., and A.M.N., Grant 926521), the National Cancer Institute (R.C.F., U2C CA252981; A.M.N., R01CA255450), the Virginia and D. K. Ludwig Fund for Cancer Research (A.M.N.), and the Donald E. and Delia B. Baxter Foundation (A.M.N.). A.M.N. is a Chan Zuckerberg Biohub – San Francisco Investigator.

## Methods

All methods are provided in the supplement.

