## Supplementary Materials for "Mapping single-cell developmental potential in health and disease with interpretable deep learning"

##### Supplementary Figures

Figure S1 CytoTRACE 2 architecture and implementation details
Figure S2 Validation and robustness of CytoTRACE 2
Figure S3 Additional benchmarking analysis and impact of dataset integration on performance
Figure S4 Extended analysis of potency programs
Figure S5 Extended analysis of developmental signatures in cancer

#### Methods

##### 1 Tumor tissue dissociation

Viably cryopreserved melanoma tumor tissue samples (*n* = 4) were thawed in a 37°C water bath and then mechanically and enzymatically dissociated. Additionally, one cutaneous metastatic melanoma tumor sample (4 mm punch biopsy) was collected in DMEM and processed within 1 hour of collection using the Miltenyi Human Tumor Dissociation Kit per the manufacturer's guidelines.

##### 2 Single-cell RNA-seq library preparation and sequencing

Single-cell-dissociated tumor tissue samples (prepared as described above) were resuspended in PBS + 0.04% BSA. Cellular viability was assessed using a hemocytometer and trypan blue with the concentration of viable cells adjusted to 1,000 cells/μl. cDNA libraries were then prepared as recommended by the 10x Genomics v2 user guide with appropriate modifications to the PCR cycles based on cDNA concentration and sequenced on an Illumina NovaSeq S4 flow cell at the Genome Technology Access Center at Washington University in St. Louis. A median sequencing depth of 50,000 reads/cell was targeted for each sample.

##### 3 The CytoTRACE 2 framework

Existing RNA-based surrogates of cellular differentiation status have notable limitations for imputing absolute differentiation states and potency categories from scRNA-seq data. For example, the original CytoTRACE, termed CytoTRACE 1 in this work, employs gene counts as an unbiased strategy for identifying immature cells[1]. Despite the utility of this approach, gene counts are subject to dataset-specific biases, making them suboptimal for potency assessment. Measures based on transcriptional entropy and RNA velocity also suffer from dataset-specific biases, a non-specific relationship to absolute differentiation status, or the requirement for continuous developmental processes within a narrowly defined time window[2-6].

Supervised machine learning (ML) models offer a potentially robust alternative to the abovementioned strategies when adequate training data are available. However, ML methods also face key challenges when applied to scRNA-seq data, including sparsity, high dimensionality, and significant data heterogeneity encompassing both biological and technical variation. While deep learning is a promising subtype of ML, often achieving remarkable performance gains over other ML methods – especially in the presence of significant complexity, noise, and uncertainty – most existing architectures lack inherent interpretability, limiting their broad applicability.

To address these challenges, we designed a novel deep learning framework that can handle the complexities of single-cell potency assessment while achieving direct biological interpretability. Unlike recent unsupervised methods[7,8] that decompose single-cell expression data into a combination of previously known and simultaneously learned new gene programs, our approach, termed a Gene Set Binary Network (GSBN), is anchored to known phenotypic states but not known gene sets. As such, GSBNs have the flexibility to discover and recover new gene programs for known phenotypic states, such as potency categories, from scRNA-seq data. As part of their design, GSBNs are highly robust and fully interpretable, meaning they can be directly interrogated to extract meaningful markers for each phenotypic class of interest across datasets, platforms, and tissues.

#### 3.1 Technical description

CytoTRACE 2 consists of five high-level components, schematically depicted in **Figures 1C** and **S1A** and described in detail below.

- *Preprocessing:* Ortholog mapping and rank-based normalization.
- *Gene set binary networks:* Identification of interpretable potency-associated gene sets for each potency category.
- *Enrichment assessment:* Evaluation of gene set activation levels in single cells.
- *Integration of scores:* Integration of gene set activation levels, both within and across gene set binary networks.
- *Postprocessing:* Leveraging transcriptional covariance and uncertainty in model predictions to smooth single-cell potency scores and produce the final output.

##### *3.1.1 Core model architecture*

Among these five components, *Gene set binary networks*, *Enrichment assessment*, *and Integration of scores* constitute the *CytoTRACE 2 core model*, a neural network architecture consisting of a shared input layer; a set of $G$ gene set binary network (GSBN) modules, where $G$ denotes the number of potency categories; and a shared output layer (**Figure S1A**). Within the core model, each GSBN module is trained to discriminate a single potency category and contains (i) a binary neural network (BNN) component, which encodes potency-associated gene sets, and (ii) downstream functions to calculate and integrate gene set enrichment scores (**Figures 1C** and **S1A**). Importantly, because weights in BNNs are constrained to binary rather than continuous values, BNNs also allow for more efficient computation and provide an implicit form of model regularization[9].

##### *3.1.2 Preprocessing*

Let input scRNA-seq dataset $\mathbf{X}$ be an $I \times C$ gene expression matrix over $I$ genes and $C$ cells. The following two pre-processing steps prepare the input dataset for training or prediction.

First, gene symbols in $\mathbf{X}$ are mapped and filtered using dictionary $\mathbb{D}$, a collection of gene symbols that harmonizes all HGNC (human) and MGI (mouse) identifiers supported by CytoTRACE 2 ("*Dictionary of input genes*" below). Following this step, the resulting expression matrix, denoted $\mathbf{X}'$, consists of $N = 14{,}271$ genes and $C$ cells. As part of this process, any genes in $\mathbf{X}'$ not present in $\mathbf{X}$ through mapping are set to zero. In the second step, $\mathbf{X}'$ is converted to rank space, yielding an $N \times C$ matrix $\mathbf{R}$, with the genes of each single-cell transcriptome $\mathbf{X}'_{*,c}$ assigned relative integer rank such that rank $1$ corresponds to the gene with highest expression. Rank space circumvents batch effects, mitigates the influence of extreme values and outliers, and reduces the risk of model overfitting. $\mathbf{R}$ is subsequently passed to the CytoTRACE 2 core model where it constitutes the model *input layer*.

##### *3.1.3 Gene set binary networks*

Input $\mathbf{R}$ is passed to each of $G$ gene set binary network (GSBN) modules within the CytoTRACE 2 core model. These modules begin by trimming $\mathbf{R}$ (**Figure S1A**) to learnable maximum rank $\tau \in \mathbb{N}$, yielding $N \times C$ matrix $\mathbf{T}$:

$$\mathbf{T}_{i,k} = \min(\mathbf{R}_{i,k}, \tau).$$

This rank trimming (see also "*Model initialization and updates*") enables calculation of enrichment scores, described in "*Enrichment assessment*" below.

Next, within each GSBN module, $M$ gene sets are learned in binary $N \times M$ matrix $\mathbf{W}^{\mathrm{B}}$, where $M \in \mathbb{N}$ is prespecified and all entries $\mathbf{W}^{\mathrm{B}}_{i,j} \in \{0,1\}$. $\mathbf{W}^{\mathrm{B}}$ constitutes the *gene set selection layer* of the CytoTRACE 2 core model; it has a continuous equivalent $\mathbf{W}$ used for model initialization and backpropagation (see also "*Core model training*"). At each forward iteration for model training, $\mathbf{W}$ undergoes binarization:

$$\mathbf{W}^{\mathrm{B}} = \mathrm{binarize}(\mathbf{W}, 0),$$

where $\mathrm{binarize}$ denotes the following utility function:

$$\mathrm{binarize}(\mathbf{M}, a)_{i,j} = \begin{cases} 1, & \mathbf{M}_{i,j} > a \\ 0, & \mathbf{M}_{i,j} \leq a. \end{cases}$$

##### *3.1.4 Enrichment assessment*

To quantify the enrichment of each gene set in $\mathbf{W}^{\mathrm{B}}$, CytoTRACE 2 leverages two complementary measures: expression enrichment score ($\mathrm{Score}_E$) and gene count enrichment score ($\mathrm{Score}_G$). $\mathrm{Score}_E$ aggregates overall expression activity of a given gene set $j$ in rank space whereas $\mathrm{Score}_G$ compares the number of expressed genes in $j$ versus background expectation, subject to $\mathbf{T}$. By integrating both scores, each providing a different axis of information, CytoTRACE 2 can learn more complex expression patterns while also achieving additional regularization through enrichment score competition. The two scores are defined as follows.

$\mathrm{Score}_E$ implements the commonly used non-parametric UCell score[10] with explicit incorporation of $\mathbf{W}^{\mathrm{B}}$. For each cell $1 \leq k \leq C$ and module gene set $1 \leq j \leq M$,

$$\mathrm{Score}_E\left(\mathbf{T}, \mathbf{W}^{\mathrm{B}}\right)_{k,j} = 1 + \left[\frac{\vec{\mathrm{S}}_j(\vec{\mathrm{S}}_j+1) - 2\sum_{i=1}^{N} \mathbf{T}_{i,k}\mathbf{W}^{\mathrm{B}}_{i,j}}{2\vec{\mathrm{S}}_j \cdot \tau}\right],$$

where $\vec{\mathrm{S}}$ denotes the vector of length $M$ containing the number of genes per gene set assigned nonzero weight in the binary weighting matrix:

$$\vec{\mathrm{S}}_j = \sum_{i=1}^{N} \mathbf{W}^{\mathrm{B}}_{i,j} \quad 1 \leq j \leq M.$$

Unlike $\mathrm{Score}_E$, $\mathrm{Score}_G$ compares the number of genes with expression rank above cutoff $\tau$ within a given gene set against the number of genes expressed in a random gene set of the same size. The latter, denoted $\widehat{\mathbf{G}}$, is estimated for gene set $j$ and cell $k$ as:

$$\widehat{\mathbf{G}}_{k,j} = \frac{\vec{\mathrm{S}}_j}{N}\sum_{i=1}^{N} \mathbf{T}^{B}_{i,k},$$

where:

$$\mathbf{T}^{\mathrm{B}} = \mathrm{binarize}(\mathbf{T}, \tau).$$

Here, $\widehat{\mathbf{G}}_{k,j}$ is the size of gene set $j$ multiplied by the background frequency of expressed genes, which is the number of genes in $\mathbf{T}^{B}_{*,k}$ normalized by $N$. Then for each cell $1 \leq k \leq C$ and module gene set $1 \leq j \leq M$,

$$\mathrm{Score}_G\left(\mathbf{T}^{\mathrm{B}}, \mathbf{W}^{\mathrm{B}}\right)_{k,j} = \frac{\sum_{i=1}^{N} \mathbf{T}^{\mathrm{B}}_{i,k} \mathbf{W}^{\mathrm{B}}_{i,j}}{\widehat{\mathbf{G}}_{k,j}}.$$

The two resulting enrichment score matrices are subsequently concatenated into a single $C \times 2M$ matrix $\mathbf{K}$:

$$\mathbf{K} = \left[\mathrm{Score}_E\left(\mathbf{T}, \mathbf{W}^{\mathrm{B}}\right) \;\; \mathrm{Score}_G\left(\mathbf{T}^{\mathrm{B}}, \mathbf{W}^{\mathrm{B}}\right)\right].$$

To transfer these enrichment scores into comparable spaces, CytoTRACE 2 standardizes each score across cells, yielding $C \times 2M$ matrix $\mathbf{K}^{\mathrm{norm}}$.

##### 3.1.5 Integration of scores

To convert the gene set enrichment scores to a single score per cell per GSBN module, the normalized scores $\mathbf{K}^{\mathrm{norm}}$ are passed through a feedforward layer, termed the *enrichment layer* in the CytoTRACE 2 core model, containing the associated length $2M$ gene set enrichment score weight vector $\vec{\mathrm{V}}$ and yielding length $C$ potency category score vector $\vec{q}$. As part of this process, dropout is applied to reduce overfitting during model training, with a predetermined fraction of the normalized scores set at random to zero. From the weights in each $\vec{\mathrm{V}}$, concatenated across potency categories into matrix $\mathbf{V}$, the directionality and importance of each gene set can be interpreted (see "*Interpretability*" below).

The model then integrates across the potency category scores produced by each GSBN module, concatenating the potency category score vectors into $C \times G$ potency score matrix $\mathbf{Q}$. This procedure represents the *shared output layer* of the CytoTRACE 2 core model.

To convert the logit entries of $\mathbf{Q}$ to likelihoods, the model applies a softmax activation function, yielding $C \times G$ matrix $\mathbf{P}$ representing the likelihood of each cell belonging to each of the six potency categories. The model then predicts cellular potency by assigning the potency category with highest likelihood for each cell, yielding length $C$ vector $\widehat{\boldsymbol{y}}$:

$$\widehat{\boldsymbol{y}}_k = \mathrm{argmax}_{\{p\}_{p=1}^{G}}\left(\mathbf{P}_{k,*}\right)$$

The $\widehat{\boldsymbol{y}}$ vector represents one of the key outputs of the CytoTRACE 2 core model. However, the model also computes an absolute developmental potential from this set of likelihoods, termed the raw potency score $\overrightarrow{\mathrm{RPS}}$. For this aspect, we introduce length $G$ ordered vector $\vec{t}$ to be multiplied by the potency category likelihood matrix:

$$\overrightarrow{\mathrm{RPS}} = \mathbf{P}\vec{t},$$
$$\vec{t} = [0.0, 0.2, 0.4, 0.6, 0.8, 1.0],$$

where $\overrightarrow{\mathrm{RPS}}$ is the length $C$ raw potency score vector. As the potency categories are ordered based on their absolute developmental potential, the resulting raw potency score will be closer to one for higher potency categories, such as totipotent, and closer to zero for lower potency categories, such as differentiated. As $\overrightarrow{\text{RPS}}$ directly incorporates model uncertainty, it is passed to "*Postprocessing*" below in order to define a more granular developmental ordering.

##### *3.1.6 Postprocessing*

Since the fully trained CytoTRACE 2 model predicts potency for each cell individually, CytoTRACE 2 further processes the output (raw potency score $\overrightarrow{\text{RPS}}$ and predicted potency categories $\hat{\boldsymbol{y}}$) to incorporate the neighborhood structure of transcriptionally similar cells. We reasoned that doing so could further improve performance given our prior experience combining gene counts with transcriptional covariance in CytoTRACE 1[1]. To this end, we devised and validated a three-step procedure using the training cohort, as described below. Importantly, this procedure improves correlations with relative developmental orderings (see "*Metrics*" below) over $\overrightarrow{\text{RPS}}$ or $\hat{\boldsymbol{y}}$ alone without sacrificing the potency classification performance achieved by $\hat{\boldsymbol{y}}$ (**Figure S1C**).

In the first step, CytoTRACE 2 applies Markov diffusion to smooth $\overrightarrow{\text{RPS}}$ using the same implementation as CytoTRACE 1[1]. In brief, the rank space gene expression input $\mathbf{R}$ is used to create a Markov matrix from the transcriptional similarity between cells over the top 1,000 genes with highest dispersion[1]. This similarity matrix is then used to smooth $\overrightarrow{\text{RPS}}$ with diffusion parameter $\alpha$ = 0.9 as previously described[1], yielding smoothed potency score $\overrightarrow{\text{SPS}}$. Using the same sampling procedure described in our previous work[1], the running time of this step can be significantly reduced without loss of performance (**Figure S1D**). In this study, sampling was restricted to datasets with >10k cells.

To reconcile $\overrightarrow{\text{SPS}}$ with predicted potency categories $\hat{\boldsymbol{y}}$, in the second step CytoTRACE 2 performs a binning procedure to maintain $\hat{\boldsymbol{y}}$ while preserving relative potency ordering within each category. To do so, CytoTRACE 2 first separates cells by their predicted potency category and assigns each cell $1 \leq w \leq C$ a rank $\mathcal{R}(k, \hat{\boldsymbol{y}}_w)$ relative to all cells sharing predicted potency category $\hat{\boldsymbol{y}}_w$. For this transformation, within each potency category $1 \leq p \leq G$, the cell with lowest potency score receives rank $1$ while the cell with highest potency score receives maximum rank $r_{\max}(p)$. Cells are then arranged uniformly by rank per potency category within equal length partitions of the unit interval, yielding binned smooth potency score $\overrightarrow{\text{SPS}}^B$. Thus, the binned smooth potency score for differentiated cells extends from 0 to 1/6, unipotent from 1/6 to 2/6, and so on, with relative ordering within each bin matching that of the original smoothed potency score.

In the third step, to further smooth $\overrightarrow{\text{SPS}}^B$ while minimizing the impact on $\hat{\boldsymbol{y}}$, CytoTRACE 2 applies k-nearest neighbor smoothing to datasets with >100 cells. To do this, $\mathbf{R}$ is first subsetted to the top 1,000 genes with highest dispersion (same as above) and passed to the ScaleData and RunPCA functions in Seurat (v4.3.0) with number of PCs set to $\min(C-1, 200)$. Next, $K-1$ nearest neighbors of each cell are identified by running nn2 from the RANN package (v2.6.1) in R (with default parameters) on the first $\beta$ principal components, where $\beta$ is ≥2 and captures $\geq$50% of the variance. $\beta$ is subsequently set to $\min(\beta, 200)$. To keep $K$ small relative to the number of cells $C$, we defined $K$ as 0.5% of $C$, with a minimum value of 3 and maximum value of 30. The $\overrightarrow{\text{SPS}}_j^B$ score for each cell $w$ is then replaced by the median $\overrightarrow{\text{SPS}}^B$ score from among $w$ and its $K-1$ nearest neighbors, yielding $\overrightarrow{\text{cytotrace2}}$, a vector of length $C$, containing the final potency scores. Categorical potency predictions are updated based on the defined intervals above, yielding $\hat{\boldsymbol{y}}^*$. We found empirically that combining these three approaches yielded superior performance on the training cohort (**Figure S1C**).

#### 3.2 Training and hyperparameter tuning

##### 3.2.1 Loss function

For model training, we defined a loss function combining cross-entropy loss with an additional term penalizing gene set size based on the binary weighting matrix $\mathbf{W}_p^{\mathrm{B}}$ originating from each GSBN module, $1 \leq p \leq G$. More precisely, given potency category predictions $\hat{\boldsymbol{y}}$ and ground truth potency categories $\boldsymbol{y}$ (see "*Gold standard scRNA-seq datasets*" below), we defined loss in terms of function $J$ given by

$$J(\hat{\boldsymbol{y}}, \boldsymbol{y}, \mathbf{W}_1^{\mathbf{B}}, \cdots, \mathbf{W}_{\mathrm{G}}^{\mathbf{B}}) = CE(\hat{\boldsymbol{y}}, \boldsymbol{y}) + \lambda \sum_{p=1}^{G} \left|(\mathbf{W}_p^{\mathrm{B}})(\mathbf{W}_p^{\mathrm{B}})^T\right|_F,$$

where $CE(\hat{\boldsymbol{y}}, \boldsymbol{y})$ denotes the cross-entropy loss, $|\cdot|_F$ denotes the Frobenius norm, and $\lambda$ denotes the gene set size penalty weight. The second term of the function $J$ serves to minimize the number of genes in each gene set while regularizing the training of the model.

To adjust for imbalances in phenotype representation, cells were partitioned by ground truth potency category and tissue source, with the loss for each cell partition computed separately first before being combined into an overall loss. Let $\mathcal{D}$ denote the set of potency category/tissue source pairs, and $\hat{\boldsymbol{y}}^d$ and $\boldsymbol{y}^d$ the predictions and ground truth potency categories respectively for cells belonging to group/tissue source pair $d \in \mathcal{D}$. Then the overall model loss is given by

$$\mathrm{Loss}(\hat{\boldsymbol{y}}, \boldsymbol{y}, \mathbf{W}_1^{\mathbf{B}}, \cdots, \mathbf{W}_{\mathrm{G}}^{\mathbf{B}}) = \frac{1}{|\mathcal{D}|} \sum_{d \in \mathcal{D}} J(\hat{\boldsymbol{y}}^d, \boldsymbol{y}^d, \mathbf{W}_1^{\mathbf{B}}, \cdots, \mathbf{W}_{\mathrm{G}}^{\mathbf{B}}).$$

##### 3.2.2 Model regularization

To promote model generalizability, we introduced two explicit regularization aspects. We included a dropout layer to avoid model overfitting to specific enrichment scores ("*Integration of scores*"). A dropout layer[11] randomly drops (sets to zero) units in a hidden layer of a neural network. This layer was applied to the normalized scores $\mathbf{K}^{\mathrm{norm}}$ during training only. Additionally, a penalty term was added to the loss function in order to constrain the number of genes in each gene set of $\mathbf{W}^{\mathrm{B}}$ ("*Loss function*").

##### 3.2.3 Model initialization and updates

Model weights were initialized according to PyTorch v2.0.0 default with the exception of the binary weighting matrices, which were initialized at random with values sampled from the Gaussian distribution with mean $-0.1$ and unit variance in order to produce a sparser initial binarization.

Model training was performed with mini-batch learning using a batch size of 1024. To balance batches and ensure equal representation for the model learning process, each batch was constructed via uniform sampling across datasets and phenotypes as implemented by torch.utils.data.WeightedRandomSampler in PyTorch.

Following initialization, forward propagation proceeded for each iteration as described in "*Core module architecture*", with parameters updated according to their definition. For numeric stability, the cutoff rank $\tau$ ("*Gene set binary networks*") for trimming input rank space expression matrix $\mathbf{R}$ was not learned directly but rather computed as a function of learnable parameter $\tau_m \in$

$\mathbb{R}$, which was initialized uniformly at random from $0 \leq \tau_m \leq 1$ per module and suitably scaled. As gene set enrichment score calculation ("*Enrichment assessment*") requires a gene set pool larger than the gene set itself for comparison, $\tau$ was computed from $\tau_m$ in such a way as to ensure that the ranks of at least 10 more genes beyond the maximum gene set size of the module were preserved following trimming to $\mathbf{T}$. Thus, at each iteration, the updated $\tau_m$ was scaled and constrained as follows:

$$\tau = 10 + \max_{1 \leq j \leq M} \vec{\mathrm{S}}_j + \mathrm{round}\big(1000 \cdot \max(0, \tau_m)\big).$$

Model predictions were assessed at each iteration against ground truth, with the loss function and its gradient computed and used to backpropagate updates to network weights using PyTorch's NAdam optimizer with custom learning rate $\mathrm{lr} = 0.001$ (see "*Hyperparameter optimization*" below) and otherwise default parameters. Given the role of inertia in successfully training binary neural networks[12,13], we employed cross-epoch gradient accumulation to dampen binary weight flipping and achieve a stabilizing effect. This approach additionally facilitates broader hyperparameter space exploration while validation-based early stopping (see *"Model evaluation and stopping"*) ensures that the most performant model encountered during training is retained. Backpropagation for the binary neural network component of each potency category module was implemented with Straight-Through Estimator and *hardtanh* activation function as previously described[9].

##### *3.2.4 Model evaluation and stopping*

We evaluated model performance over training and validation sets via category-weighted F1 score of predicted versus ground truth potency categories as implemented with *metrics.f1_score* from sklearn v1.0.2. Models were trained for 40 epochs with the best model weights by the highest score on the validation set preserved and returned for the final model.

##### *3.2.5 Hyperparameter optimization*

To evaluate the hyperparameter space of CytoTRACE 2, we performed Bayesian hyperparameter optimization over the training cohort using SigOpt (v8.8.2) (https://sigopt.com). We optimized the learning rate $\mathrm{lr}$ over $\{0.01, 0.005, 0.001, 0.0005\}$, number $M$ of gene sets per potency category over $\{1, 2, 3, 4, 5, 6, 7, 8, 9, 10, 11, 12, 13, 14, 15, 16, 32\}$, gene set size penalty weight $\lambda$ over $\{0.5, 1.0, 2.0\}$, dropout rate $\rho$ over $\{0, 0.25, 0.5\}$, and enrichment considering whether to use model expression enrichment score, model gene count enrichment score, or the concatenation of both as described in "*Enrichment assessment*" above. For every iteration of leave-one-dataset-out cross-validation, we trained models across 100 different combinations of these hyperparameters sampled based on the Bayesian optimization algorithm of SigOpt. We scored each hyperparameter combination according to the average weighted accuracy across the respective model validation sets, where weighted accuracy for each validation dataset was defined as the average of the category-weighted F1 scores per standardized phenotype and evaluated using *metrics.precision_recall_fscore_support* from sklearn v1.0.2.

We observed that variation in hyperparameter values had minimal impact on performance, underscoring overall model robustness (**Figure S1B**). Nonetheless, and consistent with expectation, combining the two gene set enrichment scores achieved superior performance compared to either enrichment score alone (**Figure S1B**, center). Moreover, when both were combined, the model was more robust to changes in other hyperparameters, such as learning rate, number of gene sets or gene set size penalty. These data supported the use of both enrichment scores rather than either enrichment score individually.

Final hyperparameter selection was done by a manual curation process identifying values yielding consistently – albeit modestly – higher weighted accuracy. In selecting the number of gene sets $M$ per potency category, we found that model performance increased with $M$ before plateauing (**Figure S1B**, right); as such, we selected $M$ slightly larger than the number corresponding to the elbow of this curve. The final hyperparameters used were $M = 12$ gene sets per potency; $\rho = 0.5$ dropout probability; $\lambda = 1$ gene set size penalty weight; and $\mathrm{lr} = 0.001$ learning rate.

##### *3.2.6 Model ensembling*

Models were trained via leave-one-dataset-out cross-validation for each of the training datasets, with final CytoTRACE 2 predictions in non-training data obtained as the result of integrating predictions across the 17 resulting models followed by an additional postprocessing step. As described in "*Integration of scores"* above, each model $m$ yields a $C \times G$ potency category likelihood matrix $\mathbf{P}^m$. Models were integrated by entry-wise averaging of potency category likelihood matrices to yield a single potency category likelihood matrix $\mathbf{P}^{ensemble}$ from which potency category predictions and raw potency scores were computed as described above, before passing them to "*Postprocessing*".

#### 3.3 Dictionary of input genes

To create dictionary $\mathbb{D}$ ("*Preprocessing*" above), all human gene symbols were mapped to their closest mouse orthologs, as determined by gene sequence similarity, using the GRCh38.p14 and GRCm39 annotation files available from Ensembl version 109, respectively. In cases where a single mouse gene $g$ was identified as the best hit for multiple human genes, the human gene with maximum sequence similarity to $g$ was selected and the remaining human gene(s) excluded from further consideration. All unique human gene symbols without orthologs by the above process were also included for completeness. To promote generalizability, only genes present in at least 80% of training datasets (with TPM/CPM > 0 in at least 1 cell per dataset) were retained. Combining these steps, $\mathbb{D}$ was assembled with 14,271 unique gene symbols, including 13,615 orthologous pairs, 521 human genes without mouse orthologs in Ensembl, and 135 mouse genes without human orthologs via the mapping step above. When mapping datasets to $\mathbb{D}$, gene symbol aliases are resolved using linked aliases available from https://biomart.genenames.org.

#### 3.4 Interpretability

The GSBN architecture of CytoTRACE 2 enables direct interrogation of the binary weight matrices, consisting of gene sets associated with each potency category (**Figures 1C** and **S1A**). By examining the orientation of the output layer weights for each gene set, we found that gene sets with positive weights (polarity) were highly enriched in a given potency category, whereas those with negative weights (polarity) were preferentially depleted (**Figure 4E**). Additionally, we reasoned that genes repeatedly selected for a given potency category were more likely to be important for effective classification. As such, we designed a metric to quantify feature importance, assigning importance scores to genes according to the frequency at which they were selected in positively versus negatively weighted gene sets. Here, we incorporate gene selection frequency across all 17 training models computed by LOOCV over the training cohort datasets.

More formally, we define $N \times G$ feature importance score matrix $\mathbf{F}$ containing the feature importance score of each gene $1 \leq i \leq N$ for each potency category $1 \leq p \leq G$ based on the gene set compositions and enrichment weights across models. Two enrichment weights correspond with each gene set, one per enrichment score type (see "*Enrichment assessment*"). Given gene set enrichment weight matrix $\mathbf{V}^l$ of model $l$, we calculate the polarity $\mathrm{Polarity}(\mathbf{V}^l, j, p)$ of gene set $j$ defined within model $l$ for potency category module $p$ as the sign of the average of these two weights. Then, relying on model binary weighting matrices to encode gene set composition, we construct feature importance score matrix $\mathbf{F}$ entry-wise as

$$\mathbf{F}_{i,p} = \sum_{l=1}^{17} \sum_{j=1}^{M} \mathbf{W}_{p,l}^{\mathrm{B}}[\mathrm{i},\mathrm{j}] \cdot \mathrm{Polarity}(\mathbf{V}^l, j, p),$$

where $\mathbf{W}_{p,l}^{\mathrm{B}}[i,j]$ denotes the $[i,j]th$ entry of the binary weighting matrix from module $p$ of model $l$.

For the analyses presented in **Figure S5A**, the top $N_p$ genes with positive feature importance score were selected for each potency category $p$. $N_p$ was defined as the median positive polarity gene set size learned by the model for each $p$.

#### 4 Gold standard scRNA-seq datasets

Classically defined potency levels, as depicted in **Figure 1A**, are not directly annotated in publicly available scRNA-seq datasets. Therefore, to train, validate, and benchmark CytoTRACE 2, we downloaded and curated 31 human and mouse scRNA-seq datasets[14-39] from peer-reviewed studies with experimentally confirmed developmental states and assignable potency levels. As part of this selection process, we applied the following inclusion and exclusion criteria to enhance experimental rigor:

- Only functionally validated developmental states supported by lineage tracing or transplantation assays were considered for analysis. Datasets with transient cell changes, such as from metabolic activation or suppression, cell cycle transitions, or environmental perturbations were excluded, as these do not represent durable developmental processes.
- Datasets with irreconcilable technical batches resulting in major imbalances in the number of cells per phenotype were excluded.
- Single-nucleus RNA-sequencing datasets were excluded, as they do not capture cytoplasmic RNA and include immature transcripts.

Among datasets satisfying these conditions, author-supplied cell type annotations were mapped to one of six standardized potency categories – totipotent, pluripotent, multipotent, oligopotent, unipotent, and differentiated – or NE (not evaluable) using established definitions ('*Potency annotations*' below). Where possible, we also examined single-cell developmental states in a dataset-specific manner and without regard to potency categories, as previously described[1]. Such 'relative' orderings, most of which were obtained from Gulati et al.[1], ranged from 1 (least differentiated) to $N$ (most differentiated) in a given dataset, and exceeded the number of resolvable potency categories in some datasets, permitting a more granular assessment.

##### 4.1 Training and test datasets

Using the abovementioned criteria, we assembled a training cohort consisting of six human and 11 mouse scRNA-seq datasets from 12 studies[14-25]. We ensured that all six potency categories were represented in both species along with a diverse array of biological (e.g., tissue types) and technical characteristics (e.g., sequencing platforms). As part of this effort, and to align with precedent in the field, we incorporated all human and mouse scRNA-seq datasets (*n* = 13) with annotatable potency categories analyzed by Gulati et al.[1] To broadly cover tissue types, we also included cell phenotypes from the Tabula Muris scRNA-seq atlas[21] for which potency categories could be determined (17 tissue types, 43 phenotypes). The resulting training cohort encompasses 175,855 cells, 18 tissue types, 88 phenotypes, and six scRNA-seq platforms (**Figure 1B**).

We next compiled a held-out test cohort that satisfies the abovementioned criteria and mirrors the training cohort with respect to species representation in each potency category. Consisting of four human and nine mouse scRNA-seq datasets from 13 studies[26-38], the test cohort spans 362,992 cells, 56 phenotypes, seven tissue types, and seven scRNA-seq platforms, including two tissue types and 21 phenotypes that were absent from training (**Figure 2C**).

To augment these data, we annotated potency categories in 459,320 evaluable cells from Tabula Sapiens, a multi-tissue scRNA-seq atlas from postmortem human donor biopsies[39]. However, given the confounding influence of postmortem intervals on human tissue mRNA levels[40], we hypothesized that Tabula Sapiens might exhibit reduced data quality. To test this, we calculated the ratio of mitochondrial (MTR) reads to total reads within each single-cell transcriptome as a proxy for overall data quality. Indeed, we calculated a mean MTR across all Tabula Sapiens tissue types, stratified by platform, of 7.4% (median of medians) – nearly 90% higher than expected for human cell types profiled by scRNA-seq data (median of medians = 3.9%; Table S1 of Osorio and Cai[41]) and 78% higher than other human datasets in the training and test cohorts, both of which include embryonic tissues with high metabolic activity (median of medians = 4.2%). Accordingly, we omitted Tabula Sapiens from the primary test cohort and evaluated it as a secondary benchmark in **Figure S3A**.

Collectively, these 31 gold standard datasets with newly annotated potency levels represent a unique community resource for systematic characterization of absolute developmental states and their molecular programs in humans and mice. Depending on platform, all scRNA-seq expression matrices were normalized to transcripts per million (TPM) or counts per million (CPM) as appropriate.

##### 4.2 Potency annotations

We used the following annotation scheme to assign single-cell transcriptomes from gold standard datasets to ground truth potency categories. Where possible, we also annotated developmental orderings within each potency category, resulting in a larger repertoire of 23 states spanning the full range of cellular ontogeny (**Figures 2A** and **2D**). However, such finer-grained orderings have sparse representation across species and scRNA-seq datasets; therefore, we restricted model training and benchmarking of absolute developmental potential to the six broader categories.

*Totipotent*. Totipotency was defined as the potential of a cell to give rise to all the cell types in a body, including extraembryonic cells, with maternal support. In addition to the totipotent zygote, we included the 2-cell stage of mouse and human embryos in which initial zygotic division has formed a 2-cell embryo. This assignment was supported by experiments in mammalian models showing development of physiologically normal organisms after re-implantation of zygote and blastomeres from the 2-cell stage[42-44].

*Pluripotent*. Pluripotency was defined as the potential of a cell to give rise to all the cell types in a body except extraembryonic cells. Although there are some reports that suggest blastomeres from the 4-cell stage are totipotent[45], given mixed reports and uncertainty in humans, we conservatively assigned the 4-cell stage as the most primitive cell state in the pluripotent category. Chronologically, the 8-cell stage and 16-cell stage – referred to as a morula – come next. The embryo then progresses to the 32-cell stage when it undergoes compaction, in which a central cavity forms and cells separate to give rise to the trophectoderm and inner cell mass, the latter of which is the origin of embryonic stem cells. As the pluripotent cells divide and specialize, they transition from the early blastocyst through the mid- and late- blastocyst stages, which mark the final stages of pluripotency in our analysis.

*Multipotent*. Multipotency was defined as the potential of an immature cell to give rise to multiple cell types (at least four) within a particular lineage. Multipotent cells exist both during early embryogenesis for specification of germ layers (mesoderm, endoderm, ectoderm) and in mature organisms as tissue-resident stem cell populations.

Within our scRNA-seq compendium, the anterior primitive streak is one of the earliest sources of multipotent cells capable of giving rise to definitive endoderm cells (e.g., epithelial cells in the respiratory and digestive tracts and the thyroid, thymus, lungs, liver, and pancreas) and mesoderm (e.g., skeletal muscle, brown fat, dorsal dermis, bone, cartilage, and smooth muscle). Using scRNA-seq and an in vitro human model of early embryogenesis, Loh et al.[17] mapped the developmental transitions from anterior primitive streak to paraxial mesoderm to somitomeres to somites to sclerotome and dermomyotome. Thus, the first tier of multipotent category consists of anterior primitive streak, while the second and the third tier consist of paraxial mesoderm and somitomere, respectively.

The fourth tier consists of early somites and pre-cranial neural crest cells (Pre-CNCCs), followed by the fifth tier of cranial neural crest cells and the sixth tier of delaminating cranial neural crest cell, dermomyotome, and sclerotome. Zalc et al.[38] performed scRNA-seq of early mouse neural crest development from pre-cranial neural crest cells (CNCCs) to CNCCs, delaminating CNCCs, and neuron/glial stem and progenitor cells and ectomesenchyme. These cells are the primitive, multilineage-producing precursors of their respective lineages as described in the original manuscripts[17,38]. To reconcile the developmental timeline of these two datasets, we anchored each within the relative embryonic time during which the cells were analyzed. As the Zalc et al.[38] pre-CNCCs were derived from the mouse 4-somite stage, the developmental timeline of CNCC fate specification started with early somites from the Loh et al.[17] study in our granular potency ordering. The post-cranial neural crest cell, like the dermomyotome, were assigned to a later multipotent stage given their shared phenotype of being early multipotent precursors to downstream tissue-resident and tissue-specific multipotent stem and progenitor cells.

The seventh tier of multipotent category consists of tissue-specific and tissue-resident multipotent stem and progenitor cells. This includes hematopoietic stem and early progenitor, intestinal stem and transit-amplifying cell, mesenchymal stem cell, multipotent pancreatic progenitor, radial glial cell, satellite stem cell. The neuron or glial stem and progenitor cell and undifferentiated ectomesenchyme (a multipotent precursor of cranial cartilage, bone, and connective tissue) were deemed equivalent to tissue-specific and tissue-resident multipotent stem and progenitor cells.

*Oligopotent*. Oligopotency was defined as the potential of an immature cell to give rise to more than one but fewer than four cell types within a particular lineage and/or cell types immediately downstream of known multipotent stem and progenitor cells. We annotated four tiers of oligopotent cells in our scRNA-seq compendium.

The first tier was labeled ‘hematopoietic stem and progenitor’, which unlike the multipotent category, is a collection of hematopoietic cells consisting mostly of downstream, specialized progenitors with a small minority of cells carrying multipotent capacity. As this group is a mixture of oligo- and very few, if any, multipotent cells, it was assigned to the oligopotent category.

The second tier consists of purified populations of cells at cellular transitions between multipotent predecessors and downstream specialized cells. For example, Ranzoni et al.[34] isolated single-cell transcriptomes from purified populations of more specialized hematopoietic progenitors, including lymphoid-myeloid progenitors (LMPs) and MK-erythroid-mast progenitor (MEMPs). Lo Giudice et al.[33] identified a cluster of cells called ‘neuroblasts’, which is the transition point between multipotent ‘progenitors’ and downstream differentiated photoreceptors, amacrine, and retinal ganglion cells. Lastly, Chan et al.[28] identified a population of cells called the ‘bone, cartilage, and stromal progenitor’, which is downstream of the multipotent skeletal stem cell but upstream of unipotent and differentiated osteogenic, chondrogenic, and stromal cells.

The third tier consists of bipotent stem and progenitor cells, including the alveolar progenitor and monocyte-dendritic cell progenitor. Alveolar progenitors are bipotent cells that give rise to type I and type II pneumocytes[46]. The monocyte-dendritic cell progenitor is a cell that gives rise to monocytes and dendritic cells[47].

The fourth tier consists of “basal” cells, a heterogeneous mixture of predominantly oligopotent progenitors with rare multipotent stem cells in each organ capable of remodeling tissue after injury. In the breast, mammary basal cells include rare bipotent stem cells, basal progenitor intermediates, and myoepithelial cells[48]. Similarly, the basal compartment in the tongue epidermis consists of a stem and bipotent progenitor cells (marked by Tp63, Sox2, Krt14, and Krt5 expression in mouse)[49]. In the skin, the basal compartment consists of stem and progenitor cell populations that give rise to the interfollicular epidermis, the hair follicle, and sebaceous gland[50]. In the proximal lung (e.g., trachea and main bronchi), the basal layer similarly consists of bipotent stem cells and downstream progenitors that generate ciliated and secretory cells[51]. Lastly, in the prostate, the basal layer consists of CD49f+Trop2+ (in human) epithelial cells that are capable of oligo-lineage regeneration[52,53]. Given that basal cells are composed of rare stem cell populations amongst mostly bipotent progenitor populations, they were classified as oligopotent, rather than unipotent or multipotent cells.

*Unipotent*. Unipotency was defined as the potential of an immature cell to predominantly give rise to one mature downstream cell type. We annotated three tiers of unipotent cells in our scRNA-seq compendium.

The first tier consists of unipotent cells with stem-like features, namely increased self-renewal capacity compared to downstream cells. This includes limbal stem cells in the cornea which self-renew and give rise to corneal epithelial cells[54,55] and nascent and adult type II pneumocytes, which can self-renew and differentiate into type I pneumocytes under certain conditions, including injury[56].

The second tier consists of unipotent progenitors with no self-renewing capacity.

The third tier consists of immature precursors cells, or transitional states between the unipotent progenitors and mature cells in a particular lineage. With the exception of CD16+CD14- 'nonclassical' monocytes, which are differentiated, monocytes derived from bone marrow were categorized as unipotent given their largely immature status and potential to give rise to downstream macrophages, while monocytes isolated from tissues were not assigned a potency category given uncertainty about their ability to generate macrophages or behave as (or give rise to) terminally differentiated monocytes (including monocytes with antigen-presenting properties)[57].

*Differentiated*. Differentiated cells were defined as mature cells within a single lineage, including terminally differentiated cells unable to make other cells, including of their same type. All cell types assigned to 'differentiated' are either mature cells or the descendants of mature and specialized cells, and most are terminally differentiated. Some mature cells within the immune system have limited potential to further differentiate or change phenotypic state when exposed to an appropriate stimulus (e.g., naïve B cells and naïve T cells can activate and further differentiate into effector B and T cells, respectively, upon antigen recognition). Monocytes isolated from blood were also classified as 'differentiated' owing to their maturity, functional specialization, phenotypic heterogeneity, and highly variable propensity to infiltrate tissue and become macrophages[57].

##### 4.3 Additional annotation considerations

For cells with identical phenotypes but different author-supplied names, we unified the annotations. For example, 'HSC-MPPs' from 'HSC development (Smart-seq2)'[34] and 'Hematopoietic stem cell progenitor (HSCP)' from 'HSPCs (C1)'[19] were annotated as 'Hematopoietic stem and early progenitor'. To balance the representation of cells from distinct lineages within a given potency category, we also re-annotated related cell subsets sharing a common parental phenotype. For example, 'CD4 helper T cells' from 'Peripheral blood (10x)'[25] and 'Tnaive/CM_CD4_activated' from 'immune cell atlas (10x)'[29] were labeled as 'T cell'. This was crucial when training CytoTRACE 2 since the probability of sampling individual cells was weighted based on phenotype. In this way, each major phenotype contributed equally during model training regardless of the number of evaluable cells, mitigating the chance of overweighting and overfitting (see "*Training and hyperparameter tuning*" above).

#### 5 Performance assessment and benchmarking

##### 5.1 Metrics

Two key metrics, each shown schematically in **Figure 1D**, were used to quantify reconstruction of known developmental orderings: absolute order and relative order. Absolute order quantifies cross-dataset performance, whereby predicted orderings from all cells with annotated potency levels are analyzed together, regardless of dataset, tissue type, or platform. Relative order quantifies performance within a given dataset and tissue type, akin to conventional pseudotime, and ranges from 1 (least differentiated) to $N$ (most differentiated) in a given dataset. For both metrics, we applied weighted Kendall correlation ($\tau$) (wdm package v0.2.4 in R) to assess concordance between known and predicted developmental orderings. Similar to our previous work[1], ground truth phenotypes corresponding to less mature cells were coded with lower ranks (starting at 1); therefore, higher predictions of developmental order were ranked such that higher values received lower ranks, and vice versa. Weighting and aggregation schemes for calculating absolute and relative order performance using Kendall correlation are described below.

- Single-cell-level performance: Absolute order and relative order performance were measured at single-cell resolution in **Figures 2E**, **S1C** bottom, **S1D**, **S2C**, **S2D**, **S2F**, **S2G**,

**S2H**, **S3A**, **S3B**, and **S3F**. Absolute order correlation was first balanced by the number of cells per dataset assigned to a given phenotype, then balanced across phenotypes within a given potency category. Relative order correlation was first balanced by the number of cells per author-supplied phenotype then balanced across standardized phenotypes (see "*Additional annotation considerations*" above) within a given relative order level.

- Phenotype-level performance: Absolute order performance was measured at phenotype-level resolution in **Figures 2A**, **2B**, **2D**, **S2E**, and **S3D**. Correlations between predicted orderings and 23 granular states were quantified in **Figures 2A**, **2D** left, and **S2E** left. To calculate this metric, predicted scores were first averaged by standardized phenotype within a given dataset $d$, then averaged across phenotypes of the same developmental state within $d$, then balanced across developmental states. Weighted Kendall correlations between predicted orderings and the six potency categories were quantified in **Figures 2B**, **2D** right, **S2E** right, and **S3D**. For this purpose, the predicted scores were first averaged by phenotype within a dataset, then weighted by number of phenotype / dataset pairs per potency category.

For categorical predictions (CytoTRACE 2 outputs only), we evaluated potency classification performance as well. Binary correctness of predicted versus ground truth potency categories was assessed via weighted accuracy (**Figure S1B and S1C**). To quantify deviations from ground truth potency on a continuous spectrum (**Figures S2A** and **S2B**), we measured the distance between each predicted potency score and the nearest subinterval or 'bin' boundary of the associated ground truth potency category (defined in step two of *"Postprocessing"* above). Statistical significance was determined by permuting CytoTRACE 2 potency scores across all evaluable cells $10^6$ times and determining the fraction of times the number of cells within 0.1 units of the correct potency bin was equal to or higher than the unperturbed number.

#### 5.2 Methods and annotated gene sets

To rigorously assess performance on our compendium of 31 curated scRNA-seq datasets, we compared CytoTRACE 2 with eight published methods for predicting developmental potential from scRNA-seq data as well as nearly 19k previously annotated gene sets. Unless otherwise stated, all evaluated methods and gene sets were applied to scRNA-seq datasets individually, without batch correction or integration across datasets, with expression data normalized per author recommendations and with default parameters. All expression data were subset to the cells with known potency. Each tissue and platform pair of Tabula Sapiens[39] and Tabula Muris[21] datasets were run separately.

Several methods rely on human gene symbols, as noted below. For all such instances, we mapped mouse dataset gene symbols to their closest human orthologs, as determined by gene sequence similarity, using the GRCm39 and GRCh38.p14 annotation files available from Ensembl, respectively. In cases where a single human gene $g$ was identified as the best hit for multiple mouse genes, the mouse gene with maximum sequence similarity to $g$ was selected.

As several methods have slower running times, to promote an equitable comparison while achieving computational feasibility, larger datasets were first downsampled. Tabula Sapiens[39] and Tabula Muris[21] datasets were each downsampled to 30 cells per phenotype, separated by tissue and platform pair, and the 'Immune cell atlas (10x)'[29], 'Human breast 1 (10x)'[18], and 'Human breast 2 (10x)'[20] datasets were downsampled to 100 cells per phenotype.

*CytoTRACE 2*. We applied CytoTRACE 2 with model ensembling and postprocessing as described in the "*The CytoTRACE 2 framework*" to predict cell potency categories and scores. Datasets containing more than 100k cells were processed in batches of 100,000 cells, and diffusion was applied in batches of 10,000 cells for datasets exceeding 10k cells. To evaluate the 17 scRNA-seq datasets included in the CytoTRACE 2 training cohort, we trained a separate model for each over the remaining 16 datasets. All other datasets were evaluated with the primary version of CytoTRACE 2 trained over all training datasets.

*CytoTRACE 1*. CytoTRACE 1, the predecessor of CytoTRACE 2, introduced transcriptional diversity quantified through gene counts as a correlate of developmental potential and exploited this concept to predict relative cellular potency from scRNA-seq[1]. CytoTRACE 1 (v0.3.3) was applied with default parameters.

*SCENT (SR)*. SCENT estimates relative cellular potency from scRNA-seq and a reference protein-protein interaction (PPI) network using single-cell signaling entropy (SR), a measure of the diversity of molecular pathway activity in a cell[3]. SCENT (v1.0.3) was executed with the "net13Jun12" human PPI network provided with the package and otherwise default parameters. For mouse datasets, genes were first mapped to human orthologs as described above. All gene symbols were converted to Entrez ID using org.Hs.eg.db (v3.15.0) in R. Gene expression matrices were normalized per documentation recommendation (https://github.com/aet21/SCENT/blob/master/vignettes/SCENT.Rmd).

*SCENT (CCAT)*. CCAT, implemented within the SCENT package, was developed as a highly efficient alternative to the original SCENT method, SCENT (SR)[2]. CCAT was applied with the same package, PPI network, and preprocessing steps described above ("*SCENT (SR)*") with expression datasets prepared per documentation recommendation.

*FitDevo*. Similar to SCENT (CCAT), FitDevo infers cellular potency from the correlation between gene expression and a measure of gene weights[58]. FitDevo (v1.2.0) was applied following tutorial instructions with binary gene weight matrix downloaded from the same source (https://github.com/jumphone/FitDevo/#demo-1---infer-developmental-potential-dp-using-expression-matrix-of-scrna-seq-data).

*SLICE*. SLICE relies on transcriptomic entropy for cellular potency prediction and lineage reconstruction, estimating entropy over functional groups of genes computed from Gene Ontology annotations[59]. SLICE (v0.99.0) was applied according to demo details from the method's GitHub page (https://github.com/xu-lab/SLICE/blob/master/demo/FB.R).

*StemID*. StemID infers cellular differentiation trajectories from scRNA-seq data with a clustering-based algorithm analyzing links between clusters[4]. StemID, implemented in RaceID (v0.1.4), was run according to documentation vignette instructions (https://cran.r-project.org/web/packages/RaceID/vignettes/RaceID.html). For each dataset, an SCseq object was initialized from each input gene expression matrix using filterData() with mintotal = 10. Ltree() and compentropy() were then applied consecutively to obtain the StemID score for cell potency.

*scTour*. scTour implements a deep learning architecture combining a variational autoencoder with a neural ordinary differential equation to reconstruct the developmental trajectory of an input scRNA-seq dataset, oriented according to gene counts[60]. scTour (v1.0.0) was trained and applied to each dataset individually per "Model training" documentation vignette instructions at https://sctour.readthedocs.io/en/latest/notebook/scTour_inference_PostInference_adjustment.ht ml. When the raw count matrix was available for the dataset, the negative binomial conditioned likelihood loss function was used. Otherwise, the CPM/TPM expression matrix was $\log_2$-transformed, and the mean squared error loss function was used instead. Cell potency scores were obtained from the developmental pseudotime predictions extracted from the model training output with get_time().

*mRNAsi*. mRNAsi utilizes a one-class logistic regression framework to construct a cellular stemness index applicable to cell potency estimation from bulk and scRNA-seq data[61]. mRNAsi was trained as described previously[1]. All input gene expression matrices were CPM/TPM normalized and $\log_2$-transformed.

*Gene sets*. The predictive capacity of 18,706 annotated gene sets (17,810 gene sets from MSigDB[62] and 896 gene sets of transcription factor binding sites from ENCODE/ChEA[63,64]) was assessed via gene set enrichment analysis. For each gene set, the AddModuleScore() function with default parameters from Seurat (v4.3.0) was applied to each expression matrix normalized via Seurat's NormalizeData() function.

#### 5.3 Comparison to scVelo

Unlike other methods assessed in this study ("*Methods and annotated gene sets*" above), scVelo[5] is limited to continuous developmental processes that occur during narrow time intervals. We therefore identified six gold standard datasets in our compendium that satisfy these requirements and span a broad range of differentiation trajectories. Loom files containing spliced, unspliced, and spanning reads for five of these datasets, namely Bone marrow (10x), Bone marrow (Smart-seq2), Peripheral glia (Smart-seq2), Dendritic cells (C1), and Intestine (Smart-seq2), were sourced from Gulati et al.[1] The sixth dataset, Pancreas (10x), was generated by Bastidas-Ponce et al.[26], but for this analysis a subsetted version was obtained from built-in datasets of the CellRank Python package[65,66] and encompassed information on the development of the murine pancreas at embryonic day 15.5 (E15.5).

For the analysis presented in **Figures S3B** to **S3D**, the *scVelo* v0.3.1 (Python package) basic velocity estimation workflow, as described in the online tutorials (https://scvelo.readthedocs.io/en/stable/getting_started.html and https://scvelo.readthedocs.io/en/stable/VelocityBasics.html), was run on the abovementioned datasets. The full dynamical model was employed to estimate velocities with default parameters for the *recover_dynamics* function. Subsequently, velocity pseudotime values were inferred using the *velocity_pseudotime* function, also with default parameters.

### 6 Robustness to variation in UMI counts and sequencing levels

To determine the influence of variable UMI counts and sequencing levels on CytoTRACE 2 (**Figure S2F**), we performed two experiments in which scRNA-seq expression data from all six droplet-based datasets in the test cohort[26,29,33,35-37] were perturbed by downsampling.

The downsampling process consisted of randomly sampling the expression data of each cell based on the transcriptome probability distribution, defined as the fractional expression of each gene after scaling the sum of UMIs in each cell to one. Using the raw count matrices, we first downsampled the expression data of each cell to the same number of UMIs: 2,000, 1,000, 750, 500, 250, 100, or 50 UMIs. Cells with UMIs lower than a given threshold were unaltered. Next, we assessed the robustness of the model to different sequencing levels by downsampling a percentage of the total UMIs per cell (90%, 75%, 50%, 25% and 10%). We randomly downsampled and repeated each analysis five times, assessed performance as described in "*Metrics*" above, and averaged the results in **Figure S2F**.

#### 7 Impact of scRNA-seq integration on performance

To validate the robustness of CytoTRACE 2 to batch effects, we assessed the concordance of CytoTRACE 2 predictions generated from two scRNA-seq datasets before and after expression integration (**Figures S3E** and **S3F**). For this purpose, we analyzed droplet and plate-seq bone marrow scRNA-seq CPM matrices from the Tabula Muris consortium[21] using fastMNN from the batchelor R package (v1.14.1). Briefly we computed 2,000 integration features for each dataset using the SelectIntegrationFeatures() function from Seurat, converted each dataset to SingleCellExperiment objects, ran the multiBatchNorm() function, and then performed batch-correction with the FastMNN function with correct.all = TRUE and otherwise default parameters.

#### 8 Analysis of mouse embryogenesis

For the analyses presented in **Figure 3**, we downloaded and curated six publicly available scRNA-seq datasets spanning each embryonic day during mouse prenatal development[15,67-72]. One dataset, which covers pre-implantation through early implantation (E0.5–E4.5) (Deng et al.[15]), was obtained from the 17-dataset training cohort and evaluated using a CytoTRACE 2 model trained on the remaining 16 datasets to avoid overfitting (see "*Methods and annotated gene sets*"). Four datasets[68-71] covering embryogenesis periods from implantation to organogenesis were previously assembled by Qiu et al. (2022)[71] and accessible through http://tome.gs.washington.edu. Finally, a recent single-nucleus RNA-seq dataset[72] covering organogenesis through birth (E8.75-P0) and generated by sci-RNA-seq3 was downloaded from http://mouse.gs.washington.edu. Since we compared CytoTRACE 2 against multiple methods with highly variable time complexity ("*Methods and annotated gene sets*"), all cells were randomly downsampled to 30 cells per author-supplied phenotype per time point, resulting in a combined dataset of 158,025 cells. This allowed us to balance considerations of performance versus computational efficiency. We ran each method on each dataset individually as described in "*Methods and annotated gene sets*" above. No dataset integration or batch normalization procedures were applied. Due to the large size of the dataset, Organogenesis (E8.75-P0)[72] was run with 10 randomly divided batches for SCENT (SR) and SLICE.

#### 9 Analysis of potency-associated molecular programs

##### 9.1 Visualization and specificity of potency programs learned by CytoTRACE 2

For the analyses presented in **Figures 4A, 4B**, **4D** (top), and **S4A**, we ran CytoTRACE 2 on each of the training and test datasets, then extracted the normalized gene set enrichment matrix $\mathbf{K}^{\text{norm}}$ (defined in "*Enrichment assessment*" above) from each of the 17 models per dataset. $\mathbf{K}^{\text{norm}}$ is derived from the penultimate layer of each GSBN module; consists of concatenated enrichment scores following batch normalization; and has a dimensionality of $C$ input cells by 2,448 when combined across the 17 models (= 17 models in the training cohort × 6 potency categories per model × 24 gene set scores per potency category [12 gene sets × 2 enrichment metrics]). We concatenated $\mathbf{K}^{\text{norm}}$ matrices from the 17 models across all 30 datasets ($C$ = 97,213 cells; downsampled as described in "*Methods and annotated gene sets*" above) to produce $\mathbf{K}_{\text{all}}^{\text{norm}}$, standardized across all cells in the training and test sets separately, and applied PCA to the resulting matrix. To adjust for differences in cell density that confound visualization, we averaged CytoTRACE 2 potency scores within each window of 1.5 PC units squared across the first two principal components. The same procedure was applied to visualize the ground truth potency of each cell (**Figure 4C**) and the predicted likelihood of each cell belonging to a particular potency category (matrix $\mathbf{P}^{ensemble}$ in "*Model ensembling*"), termed an 'enrichment index' (**Figure 4D**, top).

We assessed the specificity of potency-associated gene sets in **Figure 4D** (center and bottom) using the abovementioned enrichment index (matrix $\mathbf{P}^{ensemble}$). In brief, for a given potency category $i$, we averaged the enrichment indices for author-supplied phenotypes within each dataset, then determined the median value of the resulting quantities in each ground truth potency category. Next, we calculated the pairwise difference $\Delta_{i,j}$ between the median enrichment index of $i$ and the median enrichment index of potency category $j$ and repeated this for all $j \neq i$. We then calculated two test statistics: $\min(\Delta)$ and $\text{mean}(\Delta)$. To simulate a null distribution, we permuted the phenotype-level enrichment indices, recomputed the median enrichment index for each ground truth potency category, and calculated both statistics. We repeated this process 10,000 times. To determine an empirical p-value for potency category $i$, we tallied the proportion of times both statistics were as high (or higher) than the test statistics from the original data. We did this for each potency category in the training and test cohorts. For the statistical analyses presented in **Figures 4G** and **S4F**, we applied the same procedure but replaced the enrichment index with ssGSEA enrichment scores for the analyses in **Figures 4G** (top) and **S4F** and with $\log_2$ expression levels for the analysis in **Figure 4G** (bottom).

For the analysis presented in **Figure 4E**, we standardized $\mathbf{K}_{\text{all}}^{\text{norm}}$ across all cells within a dataset, weighted by the number of cells in each potency category. Datasets with less than two potency categories were excluded. We then aggregated the entries of $\mathbf{K}_{\text{all}}^{\text{norm}}$ across cells by mean per phenotype/dataset pair and partitioned this matrix by gene set polarity ("*Interpretability*") to form $\mathbf{K}_{+}^{\text{norm}}$ and $\mathbf{K}_{-}^{\text{norm}}$. For each resulting matrix, we then aggregated matrix columns across models and gene sets by mean per potency category, yielding a 205 × 6 matrix containing the average gene set enrichment scores per potency category for all phenotype/dataset pairs. Matrix columns were subsequently standardized to unit variance, and the matrices were transposed and visualized as heat maps.

#### 9.2 Functional annotation analysis

To interpret potency-associated genes learned by CytoTRACE 2, we applied fgsea (v1.25.1) to each rank-ordered gene list in $\mathbf{F}$ with default parameters. $\mathbf{F}$ is an $N \times G$ matrix consisting of model importance scores for all $N$ evaluable genes (=14,271) in each of $G = 6$ potency categories learned on the training cohort ("*Model interpretability*" above). Using msigdbr() (v7.5.1) in R, all mouse MSigDb signatures from HALLMARK, REACTOME, KEGG, CGP, and GO were downloaded and those with an adjusted p-value < 0.05 in at least one potency category were retained for further analysis. Due to space constraints, we selected a subset of representative molecular signatures for display in **Figure 4F**, capturing both canonical and poorly understood potency-related biology.

Among lineage restricted potency categories (multipotent through differentiated), for which established pathways have limited explanatory power, we identified molecular signatures with maximum specificity for each. To do this, we calculated the minimum delta $\Delta_{i,j}^{min}$ between the –$\log_{10}$(adjusted p-value) of a given potency category $i$ and signature $j$ and all other potency categories for signature $j$. To capture enrichment directions for signature and potency pairs with negative enrichment scores, we multiplied their corresponding –$\log_{10}$(adjusted p-value) scores by –1 prior to calculating minimum delta values. We then ranked molecular signatures in descending order of minimum delta values to identify those with maximal specificity for a given potency category. Accordingly, we identified the top molecular signatures for CytoTRACE 2 gene lists corresponding to multipotent through differentiated cells, the most specific of which are shown for multipotent and oligopotent cells in **Figures 4G** (top left) and **S4D**. The 6$^{th}$ most specific molecular signature for differentiated cells (FEVR_CTNNB1_TARGETS_UP) is highlighted in **Figure 4G** (top right) owing to its clear interpretability, marking genes upregulated by loss of Wnt/β-catenin signaling in mouse intestinal epithelium[73].

Related to **Figures S4C** to **S4E**, to decipher the molecular signatures underlying GRAESSMANN_APOPTOSIS_BY_DOXORUBICIN_DN (**Figure 4G**, top left), we ran STRING[74] (v12.0) on the corresponding leading-edge genes enriched in multipotency and downloaded enriched molecular signatures (false discovery rate < 0.05) from KEGG, REACTOME, and 'UniProt Keywords', excluding signatures related to cancer (**Figure S4C**). Protein-protein interactions among genes from fatty acid / lipid metabolism signatures (**Figure S4D**) were determined using Metascape (v3.5.20240101)[75] and visualized with Cytoscape (v3.10.1)[76]. For the analyses in **Figures S4C**, **S4E**, and **S4G**, uniformly normalized scRNA-seq data ("*Training and test datasets*" above) were first mean-aggregated by phenotype and potency within each dataset into pseudo-bulk expression profiles, then converted to rank-space across the transcriptome to mitigate technical variation. Ranked expression profiles of identical phenotypes across datasets were subsequently averaged, keeping training and test data separate. For **Figure S4C**, we averaged the ranked expression of genes within each molecular signature, then determined the mean effect size between the resulting quantities in multipotent phenotypes versus each of the remaining five potency categories. Regarding **Figure S4E**, for each gene in **Figure S4D**, we averaged the ranked expression data by potency category (keeping training and test data separate), standardized the resulting quantities by unit variance across potency categories, then median-aggregated the data across all genes for each potency category.

##### 9.3 Top potency-associated genes learned by CytoTRACE 2

For **Figure S4G**, expression data were aggregated and converted to rank space as described in "*Functional annotation analysis*" above. To identify the top CytoTRACE 2 feature with broad tissue conservation within a given potency category $p$, we first selected genes with the top 50 positive scores by potency $p$ in matrix $\mathbf{F}$ ("*Model interpretability*" above). We then calculated the area under the curve (AUC) of each of these genes in the training set as described in the caption of **Figure S4G**. Finally, we separately converted both the CytoTRACE 2 feature scores and the corresponding AUC values to rank space, averaged the resulting values, and selected the gene with the top rank from among the top 50 genes of a given $p$ in matrix $\mathbf{F}$. Such genes are referred to as 'CytoTRACE 2 informed' in **Figure S4G**.

#### 10 Candidate developmental programs in human neoplasms

##### 10.1 Survival analysis of potency markers learned from normal tissues

For the analysis presented in **Figure S5A** (left), potency-associated markers learned by CytoTRACE 2 from healthy tissues ("*Interpretability*" above) were cross-referenced with PRECOG, a database of bulk tumor gene expression profiles related to overall survival across 39 malignancies[77]. To quantify enrichment, we first mapped the mouse gene symbols from CytoTRACE 2 model to their closest human orthologs, as described in "*Methods and annotated gene sets*" above. We then rank-ordered all evaluable genes in PRECOG by their pan-cancer survival associations, summarized as meta-Z scores, and applied fgsea (v1.24.0) to each potency-associated gene set. We also repeated this process after removing genes associated with proliferation and pluripotency (**Figure S5A**, right) using published molecular signatures[62,78-81]. NES values with $Q < 0.25$ were considered significant.

##### 10.2 scRNA-seq tumor atlases

For the analyses presented in **Figures 5A** to **5E** and **S5B** to **S5H**, we downloaded 47 preprocessed scRNA-seq tumor atlases spanning 17 cancer types, 729 tumors, and 573,390 malignant cells from the Curated Cancer Cell Atlas website on June 28, 2023 (https://www.weizmann.ac.il/sites/3CA/)[82]. We selected these datasets because they encompass diverse developmental origins, including solid and hematological malignancies, and have matching cancer types in TCGA[83] and PRECOG[77], making it possible to link potency-associated molecular programs with clinical metadata at scale. For quality control, we excluded tumor samples with less than 10 malignant cells, cancer types with less than five available tumor samples, redundant samples based on duplicate identifiers, and cell line samples. We also excluded datasets with less than 10,000 genes in the CytoTRACE 2 input dictionary ("*Dictionary of input genes*" above).

In all cases, we leveraged author-supplied cell type annotations, including classifications of malignant and non-malignant cells from 3CA[82]. We also standardized non-malignant cell phenotypes based on author-provided cell types and further classified them into three broader categories based on shared lineage: immune and stromal cells in the tumor microenvironment (TME) and non-TME non-malignant cells. All non-malignant cells in each dataset were uniformly downsampled without replacement to 100 cells per standardized phenotype per tumor type for marker discovery and copy number inference. Depending on platform, all datasets were standardized to $\log_2$ CPM or TPM space as appropriate.

##### 10.3 Single-cell potency prediction and analysis

After curating scRNA-seq tumor atlases by the above protocol, we ran CytoTRACE 2 with default parameters ("*Methods and annotated gene sets*") on all annotated malignant cells from each tumor sample. We applied Seurat (v4.3.0) to visualize CytoTRACE 2 predictions for two representative tumor specimens in **Figure 5B** using NormalizeData(), FindVariableFeatures(), ScaleData(), and RunPCA(), with DimPlot() applied to the first 30 PCs. To compare CytoTRACE 2 potency scores with a proxy for overall survival associations at the single-cell level (**Figure 5B**, bottom), we selected the top 1,000 adversely prognostic genes from PRECOG[77] for each cancer type (lung adenocarcinoma, breast cancer), then used AddModuleScore() from Seurat to determine single-cell enrichment scores for each gene set.

##### 10.4 Copy number analysis

For **Figures S5D** and **S5E**, we inferred copy number profiles of individual tumor samples using inferCNV[84] (v1.12.0). As input, non-malignant cells were selected per tumor sample as described in "*Identification of potency-associated markers*". Tumors with <100 malignant cells, <10 cells in any potency category, <2 predicted potency categories, or <100 non-malignant cells were excluded. For efficiency, malignant and non-malignant cells were further downsampled to a maximum of 1,000 cells each. Following this process, we then ran inferCNV on each of the remaining 359 tumor samples with cutoff = 0.1 (droplet datasets) or cutoff = 1 (plate-seq datasets), as suggested per the online documentation, and otherwise default parameters.

To assess the relationship between predicted potency and inferred copy number patterns, we calculated a *co-association index* for each evaluable tumor sample as follows. First, we applied PCA to the Euclidean distance matrix of all single cell inferCNV profiles for a given tumor sample. Next, we applied nn2 from the RANN package (v2.6.1) in R with $K$ = 2 to the first two PCs. For cells in each potency category $p$, we tallied the number of nearest neighbors matching $p$. To determine a null distribution, we then randomized potency category labels across all single cells and repeated the above process 10,000 times, allowing us to calculate an empirical p-value for the number of nearest neighbors of $p$ with matching potency category labels expected by random chance. The minimum p-value of each tumor sample was then corrected for multiple hypothesis testing across all tumor samples using the Benjamini-Hochberg method. The resulting *Q* values are termed co-association indices.

##### 10.5 Identification of potency-associated markers

We employed the following meta-analytical procedure to identify potency-associated marker genes, prioritizing genes with consistent representation across tumor samples and datasets within a given cancer type. We began by combining, for each dataset, single-cell transcriptomes of non-malignant cells (downsampled as described above) with single-cell transcriptomes of malignant cells from each tumor sample. Following sample exclusions, only one dataset ('Glioblastoma, Neftel et al.'[85]) consisted of scRNA-seq data profiled on more than one platform; in this case, we separately combined normal cells from each platform to avoid technical biases. We then applied FindAllMarkers() in Seurat (v4.3.0) to each tumor sample separately, with only.pos = F, min.pct = -Inf, logfc.threshold = -Inf, min.cells.feature = -Inf, min.cells.group = 10, return.thresh = Inf, and otherwise default parameters. We stored the average $\log_2$ fold change (LFC) and adjusted p-value for cells of each potency category predicted by CytoTRACE 2 along with cells of non-malignant phenotypes standardized as described above ("*scRNA-seq tumor atlases*"). For non-malignant phenotypes, LFC values were further averaged within three broader phenotypic categories to balance major lineage compartments.

Next, we averaged LFC values of each phenotype (including potency categories) for tumor samples of a given cancer type $C$. This was done across datasets, filtering for common genes across datasets within cancer type $C$, yielding an $N$ by $P$ matrix $\mathbf{M}^C$, containing LFC values with $N$ genes across $P$ phenotypes. To increase the specificity of top-scoring markers, we calculated the minimum delta between $\mathbf{M}^C_{g,p}$ and $\mathbf{M}^C_{g,q}$ over all phenotypes $q \neq p$. This yielded an $N$ by $P$ matrix $\mathbf{D}^C$ containing minimum delta values for each gene $g$ and phenotype $p$ in cancer type $C$. To select the top markers, we then created a ranked gene list for each phenotype $p$ in cancer type $C$ by sorting $\mathbf{D}^C_{*,p}$ in decreasing order. This was done for all genes with $\mathbf{D}^C_{*,p} > 0.1$ and $\mathbf{M}^C_{*,p} > 0.5$. The top $T$ genes in this list were selected as potency-associated markers in cancer type $C$. If less than $T$ genes satisfied these criteria, we relaxed the $\mathbf{M}^C_{*,p}$ requirement and padded the list with the remaining genes sorted by decreasing $\mathbf{D}^C_{*,p}$ until we obtained a maximum of $T$ genes.

To select $T$, we examined the reduction in $\mathbf{M}^C_{*,p}$ as a function of $T$ across all cancer types (**Figure S5F**). Balancing considerations of detection sensitivity, as achievable by maximizing $T$, and specificity, as achievable by maximizing $\mathbf{M}^C_{*,p}$, we set $T = 20$ for all further analyses. $T = 20$ achieved 92% of the FC at $T = 10$ (when averaged across cancer types and potency categories) with twice the number of markers. To determine the statistical significance of each marker, we first converted p-values for each gene, potency category, tumor sample, and cancer type $C$ (from above) to two-sided z-scores, with directionality determined by the corresponding LFC. We then combined z-scores for each potency category across tumor samples of cancer type $C$ using Stouffer's method as previously described[77,86]. We repeated this process for unadjusted and adjusted p-values separately.

#### 10.6 Bulk tumor expression datasets

We assembled preprocessed bulk RNA-seq and microarray expression profiles from TCGA[83] and PRECOG[77] for the same 17 cancer types analyzed from 3CA. Read counts from TCGA were obtained via gdc-client v1.5.0 in June 2020. Each count matrix was normalized to TPM and $\log_2$ adjusted. Ensembl gene IDs were converted to HGNC gene symbols using the MAGeCKFlute::TransGeneID R function (v1.99.2)[87]. In total, 52 duplicate samples were removed. Clinical metadata for all samples were obtained from the GDC Data Portal (https://portal.gdc.cancer.gov/). For PRECOG, 94 bulk tumor microarray datasets and corresponding overall survival data were downloaded from https://precog.stanford.edu. Expression data in $\log_2$ space were analyzed unchanged. Datasets with a minimum expression value at least 0 and maximum >30 were $\log_2$ adjusted, then quantile normalized using normalize.quantiles() from the preprocessCore package (v1.62.0) in R.

#### 10.7 Survival analysis

To assess survival associations of predicted potency markers in TCGA and PRECOG, we first calculated the mean $\log_2$ expression of each gene set. Next, we applied Cox proportional hazards regression using coxph() from the survival package (v3.5-6) in R to calculate z-scores relating gene set expression levels to overall survival. We did this, per dataset, for tumor samples of the same cancer type. As many cancer types in PRECOG contain multiple datasets, we integrated z-scores within a cancer type using Liptak's method, with weights set to the square root of sample sizes as previously described[77]. To integrate z-scores across TCGA and PRECOG for a given cancer type, we applied Liptak's method with weights set to the square root of sample sizes, and to combine the resulting meta-z-scores across evaluable cancer types for a given potency category, we applied Stouffer's method as previously described[77,86]. Any potency categories not identifiable in a given cancer type were omitted from the meta-z-score calculation. We expressed all meta-z-scores as $-\log_{10}$ p-values with directionality for clarity.

Survival associations were calculated in various contexts using the meta-z integration scheme detailed above. For example, in **Figure 5C**, we calculated survival associations via univariable analysis, whereas in **Figure 5D**, we applied multivariable Cox regression with all evaluable potency categories included as covariates. To assess the impact of proliferation, we calculated the average expression of proliferation-related markers in each tumor using genes[62,78]. We then included proliferation as an additional covariate in bivariable Cox models incorporating each potency category. We repeated this process for pluripotency-associated markers as well[78-81]. Several analyses were restricted to TCGA owing to the availability of additional covariates. For example, we analyzed the influence of stage, age, and sex on potency-related survival associations using multivariable analysis, including pathologic stage and age as continuous variables and sex as a binary variable. We also evaluated potency-associated survival associations in TCGA after controlling for tumor grade, tumor purity (ABSOLUTE and ESTIMATE), mRNAsi[61], and TmS[88]. Precomputed mRNAsi values were obtained from Malta et al.[61] and TmS values were obtained from https://wwylab.github.io/TmS/articles/data.html.

#### 10.8 Analysis of malignant cell transcriptional hallmarks

To compare potency classifications with recently described transcriptional hallmarks in cancer[82], termed module programs, we followed the approach described at https://github.com/tiroshlab/3ca/blob/main/ITH_hallmarks/MPs_distribution/MP_distribution.R, while allowing individual cells to express >1 transcriptional program. Briefly, using datasets processed as described in "*scRNA-seq tumor atlases*", malignant cells of each tumor sample were scored for expression of each module program, for which at least 50% of genes were present in the expression matrix, using sigScores() from the scalop R package (v1.1.0). As suggested by Gavish et al., module programs with scores >1 were considered expressed (**Figure S5G**).

#### 10.9 Correlates of ICI response in melanoma scRNA-seq data

For the analysis presented in **Figure 5F**, we investigated whether candidate potency markers are associated with ICI response. To this end, we identified scRNA-seq profiles (SMART-seq2) from eight post-treatment ICI-resistant melanomas with at least 10 malignant cells each included in 3CA: 'Melanoma (Jerby-Arnon et al.)'[89]. To augment these data, we generated scRNA-seq data (10x Chromium) from five additional melanomas treated with ICIs, including three patients in whom durable clinical benefit (DCB) was achieved ("*Human patient samples*" above). Using Seurat (v4.3.0) to preprocess these newly generated samples, we first removed cells with <200 detectable genes and mitochondrial gene ratios >0.25. We then ran NormalizeData(), FindNeighbors() with dims = 1:10, FindClusters() with default parameters, and RunUMAP() with dims = 1:10. We removed clusters with >50% of cells predicted as doublets using DoubletFinder[90] (v2.0.3). We then annotated clusters based on expression of canonical marker genes (positive markers unless indicated otherwise): *MITF*, *S100B*, *MLANA*, *PMEL*, or *TYR* = melanoma cells; *EPCAM* = epithelial cells; *VWF* or *PECAM1* = endothelial cells; *COL1A1* or *COL3A1* = fibroblasts; *ACTA2* or *TAGLN* = smooth muscle cells; *CD79A* or *CD79B* = B cells; *IGKC* or *MZB1* = plasma cells; *CD3D* and *CD4* or *IL7R* = CD4 T cells; *CD3D* and *CD8A* or *CD8B* = CD8 T cells; low *CD3D* and high *GNLY* = NK cells; *CD68* or *CD163* = macrophages; *CD1C* or *CD209* = dendritic cells. Malignant and non-malignant classifications were corroborated by inferCNV, applied as above ("*Copy number analysis*").

Using melanoma-specific potency markers defined from 3CA datasets, for which markers of multipotent-like, oligopotent-like, and differentiated-like cells were defined, we first harmonized across platforms (Smart-seq2 and 10x Chromium) by retaining only those genes present in all 13 evaluable expression matrices. We then calculated the mean $\log_2$ expression, denoted $\overline{\mathrm{LE}}$, of each potency category in malignant cells from each sample. To combat technical variation across samples, we further normalized $\overline{\mathrm{LE}}$ values using a potency category with "opposite" predicted developmental potential. More specifically, within each sample, we subtracted the $\overline{\mathrm{LE}}$ of differentiated-like cells from the $\overline{\mathrm{LE}}$ values of multipotent-like and oligopotent-like cells. Separately, we averaged the $\overline{\mathrm{LE}}$ values of multipotent-like and oligopotent-like cells, then subtracted the resulting quantities from differentiated-like cells.

#### 10.10 Correlates of ICI response in bulk tumor expression datasets

For **Figure 5G**, preprocessed bulk tumor RNA-seq data and corresponding clinical information from 601 patients with cancer treated with anti-PD-1 monotherapy or anti-PD-1/anti-CTLA-4 combination therapy ICIs were obtained from public repositories, including TIGER[91] (http://tiger.canceromics.org/#/home), TIDE[92] (http://tide.dfci.harvard.edu/), MORRISON[93] (https://github.com/ParkerICI/MORRISON-1-public), https://github.com/vanallenlab/schadendorf-pd1[94], GSE166449[95], and PRJNA371283[96]. For **Figure S5I**, preprocessed bulk RNA-seq data from mouse tumor models treated with ICIs, incorporating treatment and response annotations, were downloaded from TISMO[97] (http://tismo.cistrome.org/). All datasets were either obtained or processed as $\log_2$ TPM prior to analysis, and only cancer types present in our 3CA analysis were considered.

To quantify the enrichment of candidate potency markers from each evaluable cancer type, we first standardized expression for each gene via unit variance normalization in each dataset, subsetting by ICI time point and treatment to mitigate batch effects. We then repeated the same procedure for scoring potency-associated markers as described above in "*Correlates of ICI response in melanoma scRNA-seq data*" to ensure consistency. Expression enrichments were averaged by dataset, ICI response status, and ICI treatment in **Figures 5G** and **S5I**.

#### 11 Quantification and statistical analysis

Relationships between two ordered variables were assessed by correlation tests or linear regression. Two-group comparisons were assessed using unpaired or paired Wilcoxon tests, as appropriate. Differences across three or more groups were analyzed using a Kruskal-Wallis test. Results with $P < 0.05$ were considered significant. Data analyses were performed with Python (v3.9.0) and R (v4.2.0+).

#### 12 Software implementation

R and Python packages for running CytoTRACE 2 with the pre-trained model are available at https://github.com/digitalcytometry/cytotrace2. Both packages implement optional parallel processing for efficient execution (**Figure S1D**) and built-in plotting functions (UMAPs, box plots). Documentation, vignettes, and input examples are provided.

#### 13 Data and code availability

Gold standard datasets with known developmental orderings used in this study are available from the Gene Expression Omnibus (GEO), ArrayExpress, or the Sequence Reach Archive (SRA): GSE52583 ('AT2/AT1 lineage (C1)'[23]), GSE109774 ('Bone marrow (10x)', 'Bone marrow (Smart-seq2)', and 'Tabula Muris (Smart-seq2/10x)'[21]), GSE60783 ('Dendritic cells (C1)'[22]), GSE97391 ('Direct in vitro neuron (inDrop)' and 'Standard in vitro neuron (inDrop)'[14]), GSE70245 ('HSPCs (C1)'[19]), GSE113197 ('Human breast 1 (10x)' and 'Human breast 1 (C1)'[18]), GSE161529 ('Human breast 2 (10x)'[20]), GSE36552 ('Human embryo (Tang et al.)'[24]), GSE92332 ('Intestine (Drop-seq)' and 'Intestine (Smart-seq2)'[16]), GSE85066 ('Mesoderm (C1)'[17]), GSE45719 ('Mouse embryo 1 (Tang et al.)'[15]), SRP073767 ('Peripheral blood (10x)'[25]), GSE128639 ('BM-MNC (CITE-seq)'[37]), GSE100866 ('Cord blood (CITE-seq)'[36]), E-MTAB-9067 ('HSC development (Smart-seq2)'[34]), GSE90742 ('HSCs and MPPs (inDrop)'[35]), E-MTAB-11536 ('Immune cell atlas (10x)'[29]), GSE76408 ('Lgr5-CreER intestine (CEL-seq)'[32]), E-MTAB-3321 ('Mouse embryo 2 (Smart-seq2)'[31]), GSE59892 ('Mouse embryo 3 (Smart-seq)'[27]), GSE162044 ('Neural crest (Smart-seq2)'[38]), GSE132188 ('Pancreas (10x)'[26]), GSE99933 ('Peripheral glia (Smart-seq2)'[30]), GSE122466 ('Retinal neurons (10x)'[33]), GSE64447 ('Skeletal stem cell (C1)'[28]), GSE201333 ('Tabula Sapiens (Smart-seq2/10x)'[39]), GSE67602 ('Hair Epidermis (C1)'[98]), and GSE75330 ('Oligodendrocyte phenotypes (C1)'[99]).

CytoTRACE 2 is freely available for non-profit academic use at https://github.com/digitalcytometry/cytotrace2.

#### Supplementary Figures

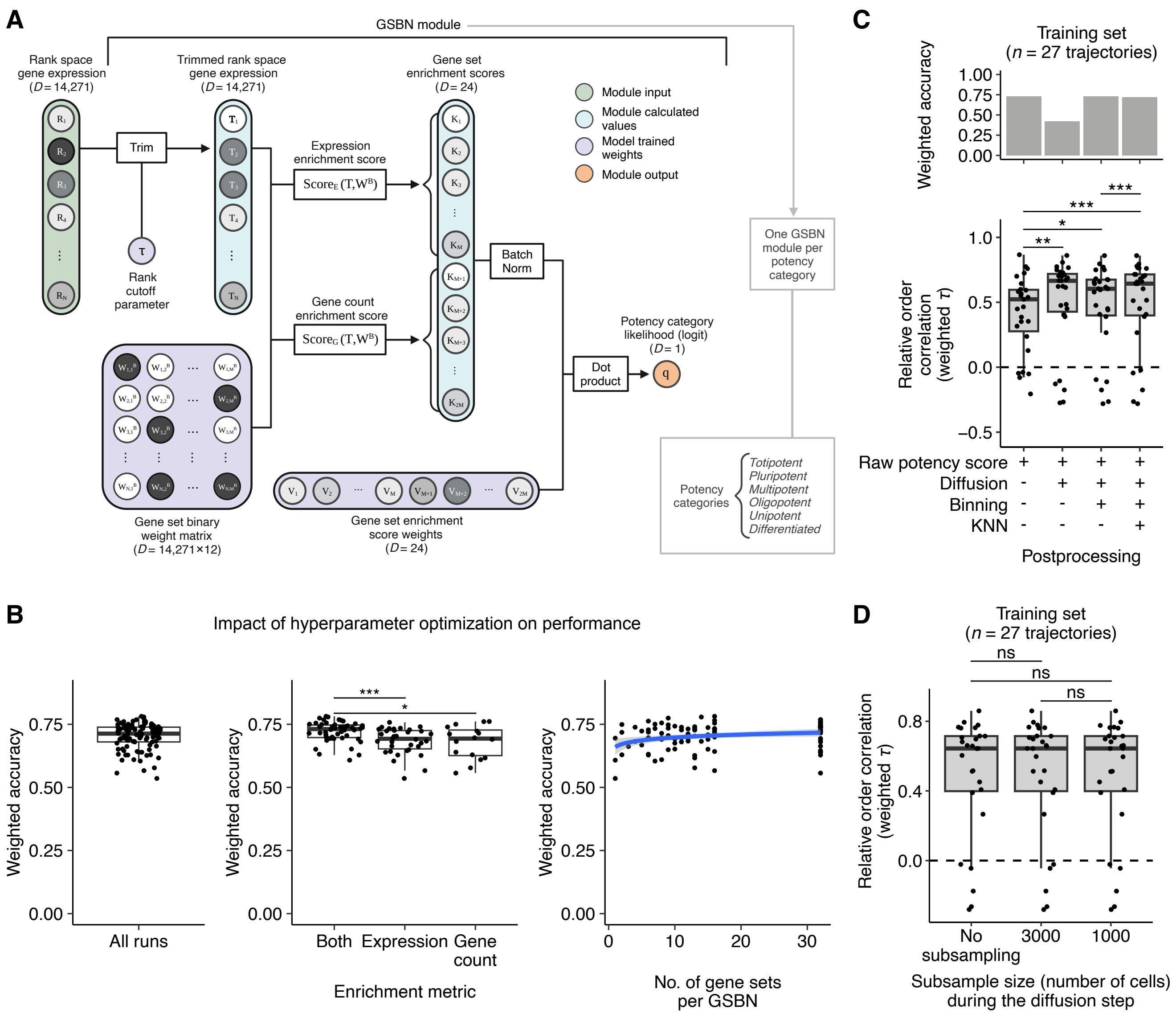

**Figure S1. CytoTRACE 2 architecture and implementation details.** (**A**) Schematic overview of the CytoTRACE 2 core model, focusing on architectural details and key operations between the input layer (a rank-ordered single-cell gene expression profile) and output layer (potency likelihood estimation) for a single gene set binary network (GSBN) module. Note that six GSBN modules, one per potency category, are included in the full model, as illustrated in **Figure 1C**. All components, variables, and operations are described in **Methods**. (**B**) Impact of hyperparameter optimization on performance. *Left*: Scatter plot showing the weighted accuracy (**Methods**) for single-cell potency prediction from the training set using LOOCV. Each point ($n$ = 100) denotes the results from one iteration of Bayesian hyperparameter optimization. *Center:* Same as the left but showing weighted accuracy as a box plot stratified by models using one or two gene set enrichment metrics: an 'expression' enrichment score and a 'gene count' enrichment score (**Methods**). Statistical significance was determined using a two-sided unpaired Wilcoxon test. *Right:* Same as the left but showing weighted accuracy plotted by the number of gene sets per GSBN module. The blue line denotes logarithmic regression, and the band indicates 95% CI. (**C**) Box plot showing the serial impact of three post-processing procedures

("*Post-processing*" in **Methods**) on the prediction of relative ordering (bottom) and weighted accuracy (top) across training datasets (denoted by points) (**Methods**). All input scores to postprocessing procedures were obtained via LOOCV. (**D**) Box plot showing the performance of CytoTRACE 2 for the prediction of relative ordering after subsampling cells for the Markov diffusion step of the postprocessing procedure ("*Postprocessing*" in **Methods**). All training datasets were analyzed using LOOCV. In B–D, the box center lines, bounds of the box, and whiskers denote medians, 1st and 3rd quartiles, and minimum and maximum values within 1.5 × IQR (interquartile range) of the box limits, respectively. Statistical significance in C and D was determined using a two-sided paired Wilcoxon test. *$P$ < 0.05; **$P$ < 0.01; ***$P$ < 0.001. Note that tissue-specific expression matrices from plate- and droplet-based platforms within Tabula Muris[21] were analyzed individually in C and D for clarity, yielding 27 total trajectories. A was created using BioRender.com.

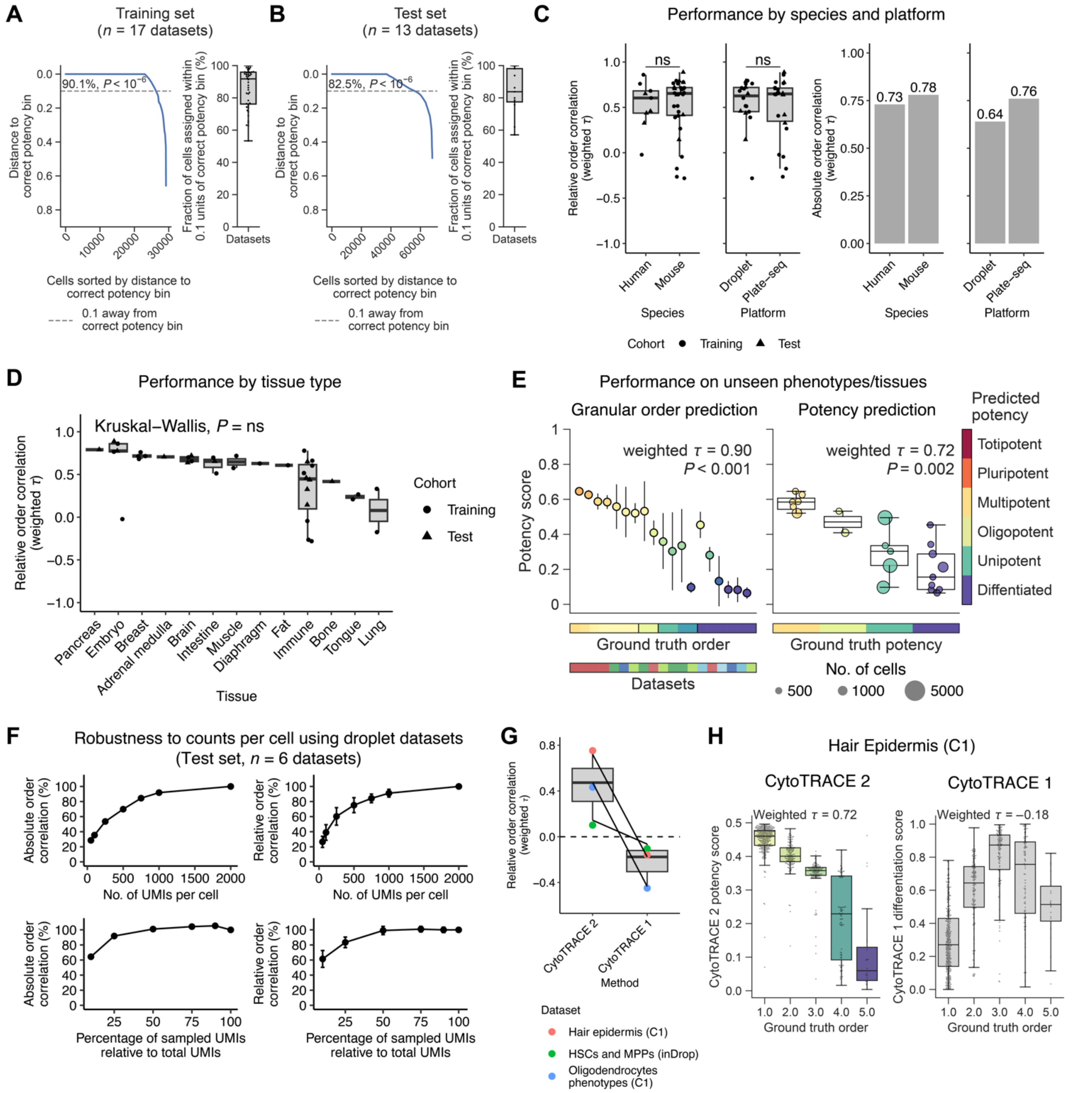

**Figure S2. Validation and robustness of CytoTRACE 2.** (**A**) *Left:* Line plot showing the minimum distance between the predicted potency score of each cell in the training set and the ground truth potency 'bin'. Potency bins are defined as ordered partitions of equal size (1/6 each) between 0 (differentiated) and 1 (totipotent), where differentiated cells occupy the bin from 0 to 1/6, unipotent cells occupy the bin from 1/6 to 2/6, and so on (**Methods**). The percentage of cells within 0.1 units of the correct bin is indicated (dashed line). Statistical significance was determined by a permutation test (**Methods**). *Right:* Box plot summarizing the fraction of cells within 0.1 units of the correct bin for each training dataset. (**B**) Same as A but for the held-out test set. (**C**) CytoTRACE 2 performance for reconstructing relative order (left) and absolute order (right) by species and platform. Performance was evaluated at the single-cell level using weighted Kendall correlation, as described in **Methods**. For relative order assessment, 40 trajectories with at least two developmental states each were evaluable. Statistical significance in the left panel was determined using a two-sided unpaired Wilcoxon test. (**D**) Same as panel C (left) but showing relative order performance stratified by tissue type, with statistical significance determined by a Kruskal-Wallis test. (**E**) Same as **Figure 2D** but showing performance on 21 cell phenotypes that were unseen during model training. (**F**) Analysis of robustness to the number of UMIs per cell (top) and the percentage of the original number of UMIs per cell (bottom) using all six droplet datasets in the test set[26,29,33,35-37], shown for absolute order (left) and mean relative order (right). Performance was analyzed using weighted Kendall correlation (**Methods**) and expressed as a percentage of the results obtained on a maximum of 2,000 UMIs per cell (top) and unperturbed data (bottom). Error bars represent S.E.M. Results for each dataset represent the average across five rounds of UMI downsampling. (**G**, **H**) Analysis of trajectories inverted by the original CytoTRACE algorithm[1] (termed CytoTRACE 1). (**G**) Paired box plot comparing relative order correlations obtained by CytoTRACE 1 and CytoTRACE 2 on three datasets inverted by CytoTRACE 1[35,98,99]. (**H**) Box plots showing CytoTRACE 2 and CytoTRACE 1 performance on a representative dataset (Hair Epidermis (C1)[98]), with relative order annotations obtained from Gulati et al.[1] Each point denotes a cell, with performance quantified by Kendall correlation weighted by the number of cells in each known developmental state. In panels A–E and G–H, the box center lines, bounds of the box, and whiskers denote medians, 1st and 3rd quartiles, and minimum and maximum values within 1.5 × IQR (interquartile range) of the box limits, respectively.

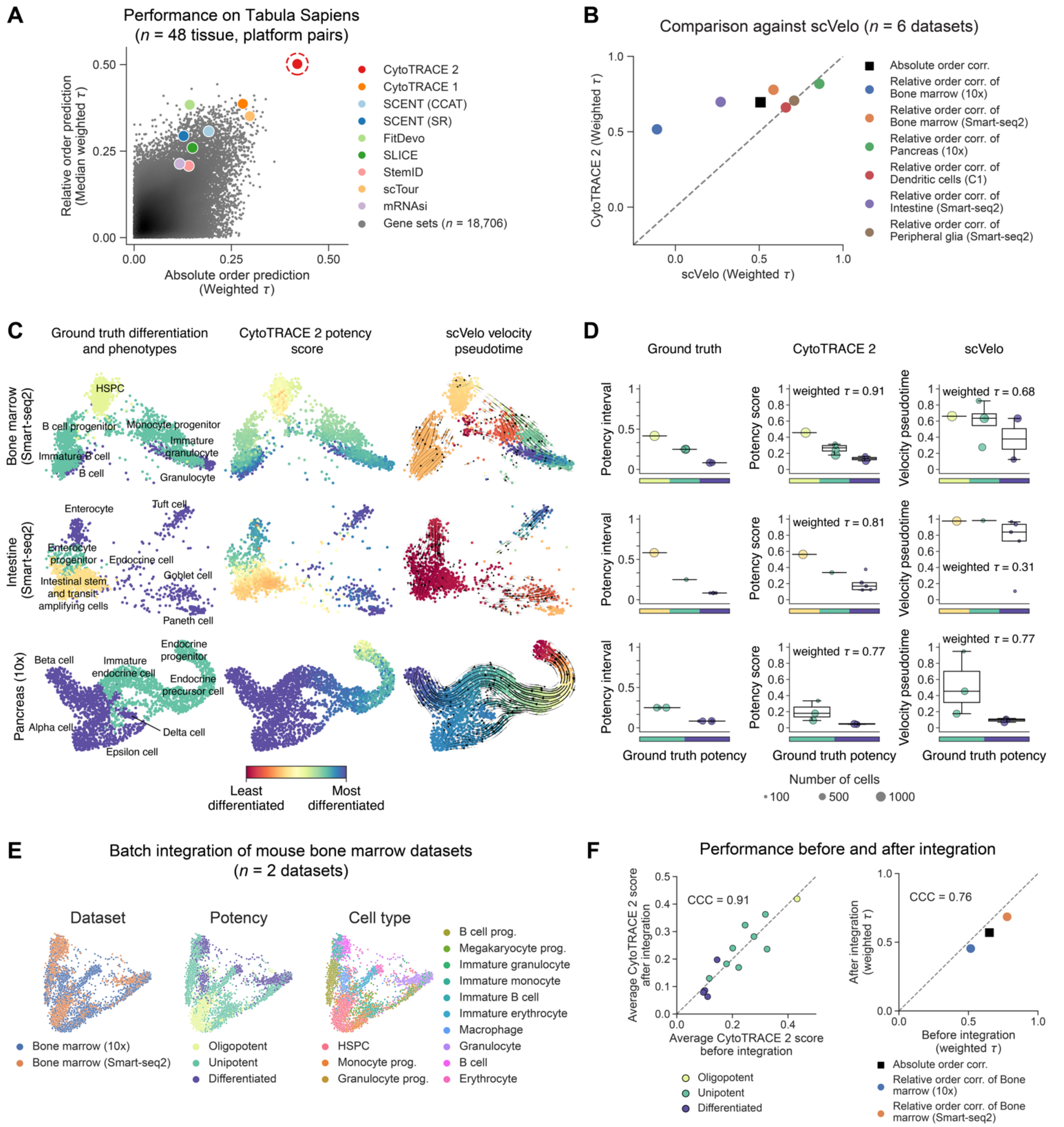

**Figure S3. Additional benchmarking analysis and impact of dataset integration on performance.** (**A**) Same as **Figure 2E** (right) but showing reconstruction of ground truth developmental orderings in Tabula Sapiens[39] across 48 tissue / platform pairs for absolute ordering assessment (x-axis) and 28 tissue / platform pairs with at least two developmental states each for relative ordering assessment (y-axis). (**B–D**) Comparative analysis of CytoTRACE 2 and scVelo. (**B**) Scatter plot comparing the absolute and relative ordering performance of CytoTRACE 2 and scVelo on six scRNA-seq datasets with continuous developmental trajectories (**Methods**). Performance was assessed at the single-cell level using weighted Kendall correlation ($\tau$) (**Methods**). (**C**) UMAP projections of three representative datasets analyzed in panel B, distinguishing ground truth potency, CytoTRACE 2 potency scores, and scVelo pseudotime predictions by color. Velocity fields are also shown for the latter. (**D**) Box plots depicting ground truth potency categories by phenotype (left) and CytoTRACE 2 and scVelo scores mean-aggregated by phenotype (center and right, respectively) for the same datasets in panel C, ordered top to bottom. Kendall correlations ($\tau$) against ground truth were weighted by the number of phenotypes per potency category. The box center lines, bounds of the box, and whiskers denote medians, 1st and 3rd quartiles, and minimum and maximum values within 1.5 × IQR (interquartile range) of the box limits, respectively. (**E**, **F**) Impact of scRNA-seq integration on CytoTRACE 2 performance. (**E**) PCA embeddings of two mouse bone marrow datasets following batch integration using FastMNN (**Methods**), colored by datasets, cell types, and ground truth potency categories. (**F**) Scatter plots comparing CytoTRACE 2 results before and after integration, showing average CytoTRACE 2 potency scores of 13 standardized phenotypes (left) and performance (right), with concordance assessed by Lin's Concordance Correlation Coefficient (CCC), which jointly measures linearity and deviation from the y=x diagonal. Absolute and relative orderings were determined at the single-cell level using weighted Kendall correlation (**Methods**).

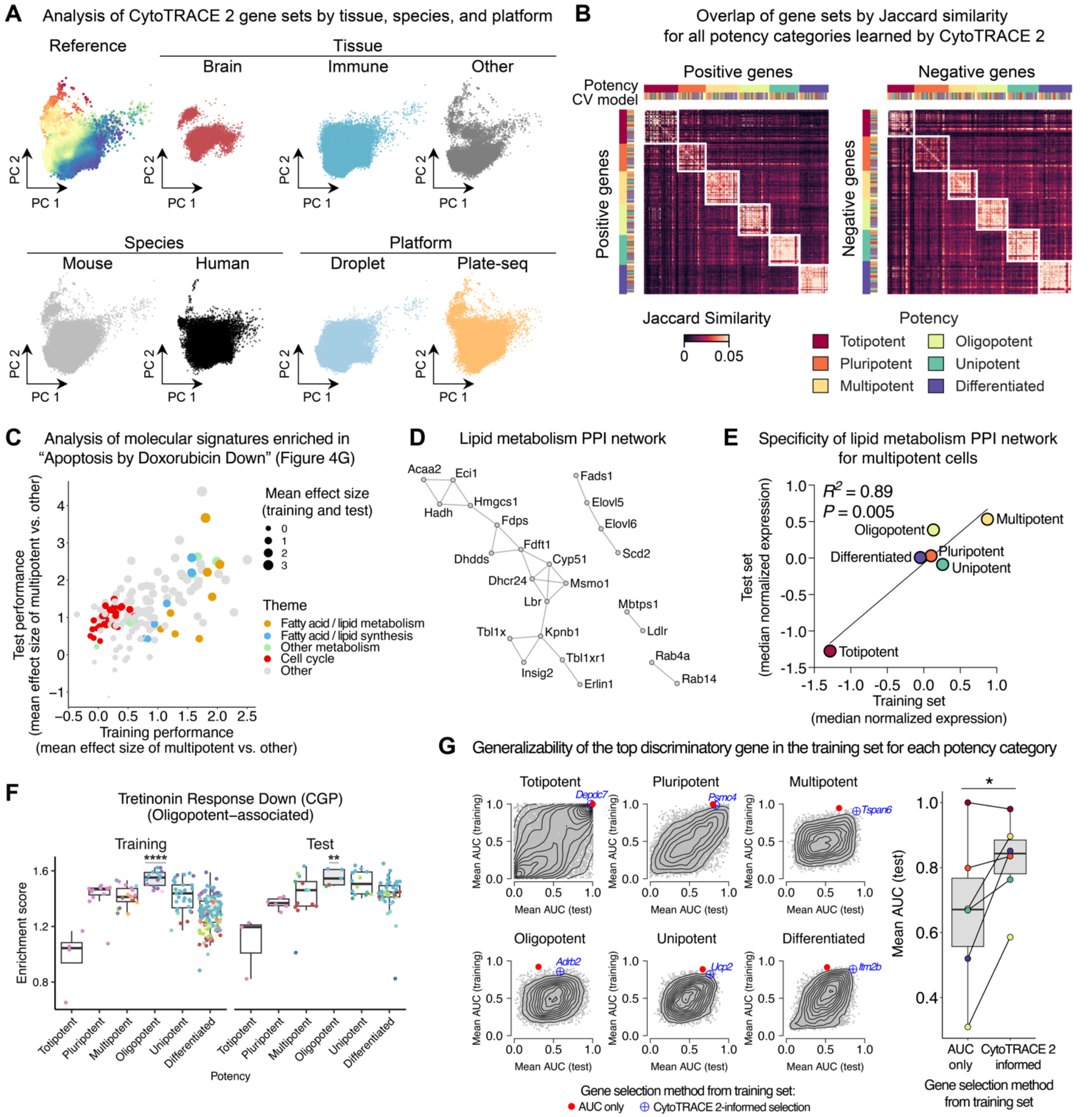

**Figure S4. Extended analysis of potency programs.** (**A**) Same as **Figure 4B** but colored by tissue type (three broad categories shown for clarity), species, and scRNA-seq platform. The embedding in **Figure 4B** is shown as a reference in the upper left. (**B**) Heat map depicting pairwise similarity of all gene sets learned by CytoTRACE 2 on the 17-dataset training set, aggregated across all 17 models. Overlap was quantified by Jaccard index and stratified into gene sets with positive (left, $n = 668$) and negative weights (right, $n = 556$); gene set polarity was determined as described in "*Interpretability*," **Methods**. (**C** – **E**) Analysis of a lipid metabolism network associated with multipotency in training and test data. (**C**) Scatter plot showing the discriminatory power of molecular signatures enriched within the GRAESSMANN_APOPTOSIS_BY_DOXORUBICIN_DN signature, a marker of multipotent cells (**Figure 4G**, top left). Discriminatory ability was assessed by calculating the mean effect size between the enrichment of each gene set within multipotent phenotypes versus enrichment in each of the remaining five potency categories (**Methods**). Gene sets are colored by dominant functional themes, with fatty acid and lipid-related processes skewed toward higher effect size and cycling-related processes skewed toward lower effect size. (**D**) Genes from fatty acid / lipid metabolism signatures in panel C that encode proteins exhibiting evidence of physical interactions (*n* = 24 genes; **Methods**). PPI, protein-protein interaction. (**E**) Scatter plot showing the mean expression of genes in panel D across all unique phenotypes, stratified by ground truth potency within training and test sets. Expression was normalized as described in **Methods**. Concordance and significance were assessed by linear regression (solid line) and a two-sided t test, respectively. (**F**) Same as **Figure 4G** (top) but showing a molecular signature (MARTENS_TRETINOIN_RESPONSE_DN[100]) with maximal specificity for oligopotent cell markers learned by CytoTRACE 2 (**Methods**). (**G**) *Left:* Scatter plots with overlaid contour lines showing the sensitivity and specificity of each gene in the CytoTRACE 2 input dictionary (*n* = 14,271; **Methods**) for distinguishing each potency category, separated by training and test datasets. For potency category $p$, genes are ranked by the mean area under the curve (AUC) between their expression in $p$ and each of the remaining five potency categories. Two genes are highlighted in each plot, the one with highest AUC in the training cohort (red circle) and the leading gene informed by CytoTRACE 2 feature weights (blue crosshair; **Methods**) in potency category $p$. Gene symbols corresponding to the latter are indicated in blue. The top CytoTRACE 2 informed genes for multipotent and differentiated cells are highlighted in **Figure 4G** (bottom). *Right:* Box plots comparing the mean AUCs in the test cohort for the two selection methods in the left panel (red circle and blue crosshair), colored by potency category (same as panel B legend). Statistical significance was determined with respect to ‘AUC only’ with a one-sided paired Wilcoxon test. For additional details, see **Methods**. In C, E, and G, to remove imbalances in cell and phenotypic representation, pseudo-bulk expression profiles of each phenotype/dataset combination were analyzed as described in **Methods**. In F and G (right), the box center lines, bounds of the box, and whiskers denote medians, 1st and 3rd quartiles, and minimum and maximum values within 1.5 × IQR (interquartile range) of the box limits, respectively. $^{**}P < 0.01$; $^{****}P < 0.0001$.

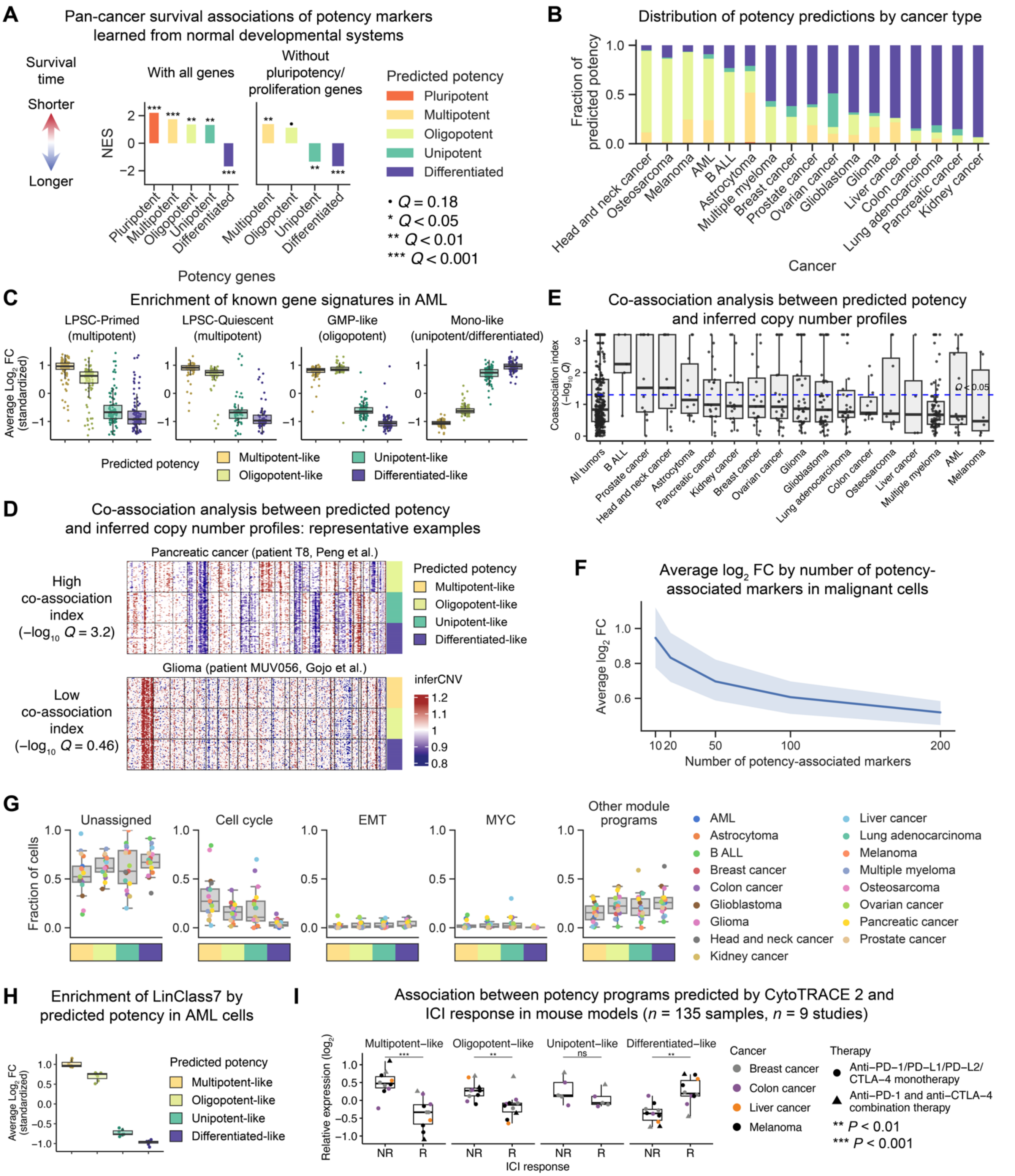

**Figure S5. Extended analysis of developmental signatures in cancer.** (**A**) *Left*: Bar plot showing enrichment scores of potency-associated genes learned by CytoTRACE 2 from healthy tissues (**Figure 4**) in 23,287 human genes ranked by their survival associations across 39 malignancies (PRECOG database[77]). Enrichments were determined by pre-ranked gene set enrichment analysis (GSEA); Normalized enrichment score (NES) values indicate relationships with overall survival: positive, adverse association; negative, favorable association. *Right*: Same as the left panel but with 2,737 genes previously linked to proliferation, pluripotent stem cells, or pluripotency-associated genes learned from CytoTRACE 2 omitted from PRECOG before running GSEA (**Methods**). (**B**) Distribution of single-cell potency predictions per cancer type, shown for all tumor specimens with scRNA-seq profiles summarized in **Figure 5A**. Colors are defined in the legend of panel A. (**C**) Box plots showing relative expression levels of cell state signatures from patients with acute myeloid leukemia (AML) in 13,420 AML cells[101] stratified by potency categories identified by CytoTRACE 2. Each point denotes a single gene from the corresponding gene set ID indicated above the plot. Genes were internally normalized within each tumor sample as the mean $\log_2$ fold change (FC) within a given potency category versus the remaining cells in the tumor, as described in "*Identification of potency-associated markers*" (**Methods**), then z-score normalized (standardized) across potency categories. The four signatures[102], LSPC-Primed, LSPC-Quiescent, GMP-like, and Mono-like, are expected to be most highly expressed in multipotent, multipotent, oligopotent, and unipotent/differentiated cells, respectively. (**D**, **E**) Co-association between CytoTRACE 2 potency predictions and inferred copy number profiles for 359 evaluable tumor samples with scRNA-seq data (**Methods**). (**D**) Representative examples of tumor samples with and without significant potency / copy number co-associations, as determined by a co-association index described in **Methods**. For clarity, individual cells were downsampled to balance representation per predicted potency category. (**E**) Box plots aggregating $-\log_{10}$ *Q*-values of co-association indices by and across tumor types. Points denote individual tumor samples. (**F**) Plot depicting the mean $\log_2$ FC of markers identified for each potency category predicted by CytoTRACE 2 from scRNA-seq profiles of malignant cells (matrix $\mathbf{M}_{*,p}^{C}$ in "*Identification of potency-associated markers*", **Methods**), aggregated across all potency categories and evaluated cancer types. The blue line and band indicate the mean and 95% CI across cancer types, respectively. (**G**) Box plots showing previously described transcriptional hallmarks ("module programs") in cancer[82] versus single-cell potency assignments predicted by CytoTRACE 2. Points denote the fraction of cells per tumor type ($n$ = 17) assigned to a given cancer hallmark (or not assigned to any hallmark), stratified by predicted potency category. Colors are defined in the legend of panel A. (**H**) Same as panel C but showing the relative expression levels of a previously defined AML drug response signature (LinClass7), comprised of seven genes, expected to be most highly expressed in immature AML cells[102]. (**I**) Same as **Figure 5G** but shown for 135 bulk RNA-seq profiles of mouse tumor models treated with ICIs. R, responder; NR, non-responder. In panels C, E, and G–I the box center lines, bounds of the box, and whiskers denote medians, 1st and 3rd quartiles, and minimum and maximum values within 1.5 × IQR (interquartile range) of the box limits, respectively.
